# A synthetic distributed genetic multi-bit counter

**DOI:** 10.1101/2021.11.10.468063

**Authors:** Tianchi Chen, M. Ali Al-Radhawi, Christopher A. Voigt, Eduardo D. Sontag

**Author notes:** These authors contributed equally (Tianchi Chen, M. Ali Al-Radhawi). This research was supported in part by Grants SRC SB-2837-B and NSF 1849588, for the project “Very large-scale genetic circuit design automation”.

## Abstract

A design for genetically-encoded counters is proposed via repressor-based circuits. An *N* -bit counter reads sequences of input pulses and displays the total number of pulses, modulo 2*^N^* .

The design is based on distributed computation, with specialized cell types allocated to specific tasks. This allows scalability and bypasses constraints on the maximal number of circuit genes per cell due to toxicity or failures due to resource limitations.

The design starts with a single-bit counter. The *N* -bit counter is then obtained by interconnecting (using diffusible chemicals) a set of *N* single-bit counters and connector modules.

An optimization framework is used to determine appropriate gate parameters and to compute bounds on admissible pulse widths and relaxation (inter-pulse) times, as well as to guide the construction of novel gates.

This work can be viewed as a step toward obtaining circuits that are capable of finite-automaton computation, in analogy to digital central processing units.

**Graphical Abstract:** 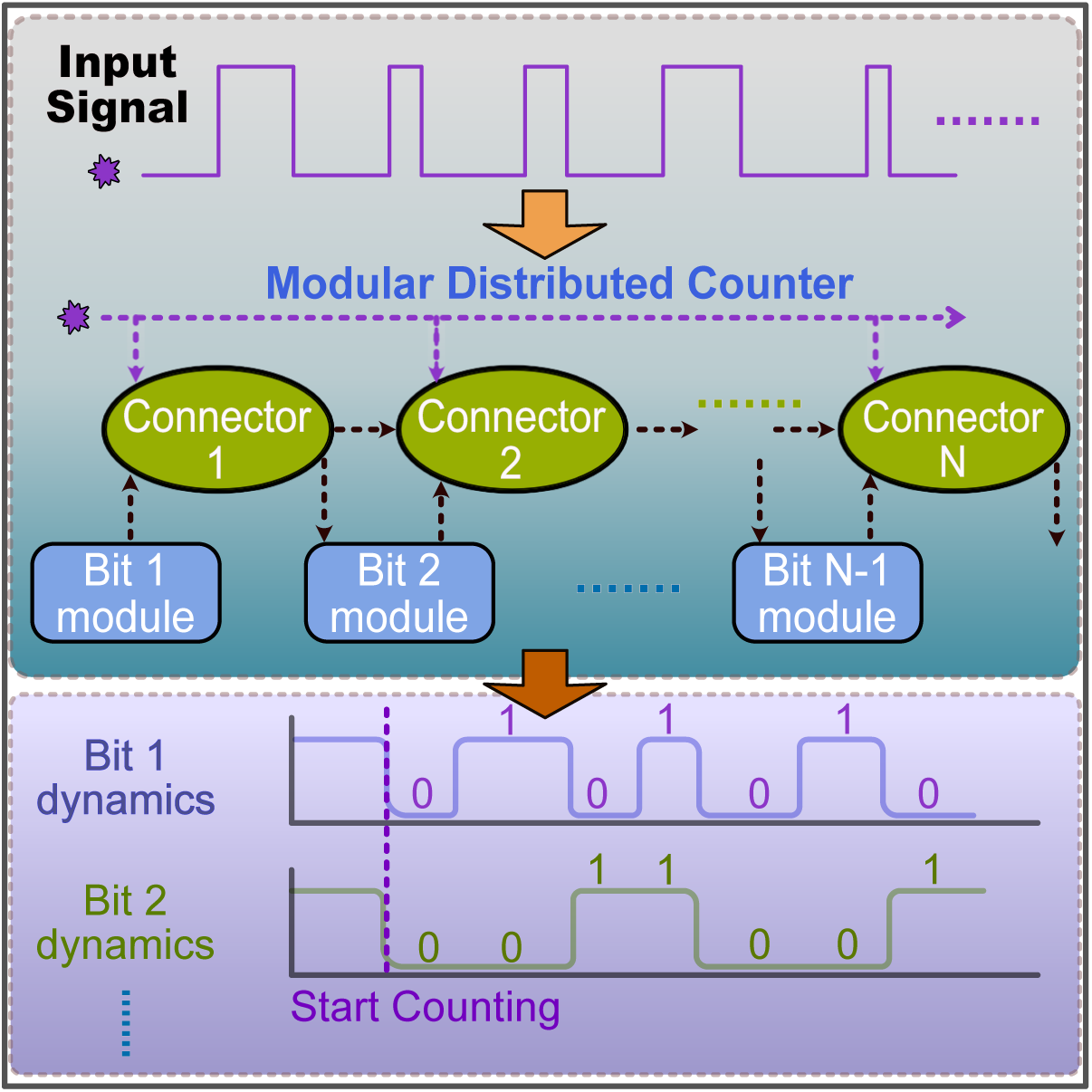

## 1 Introduction

We introduce a new design of counters: an *N* -bit counter which reads sequences of input pulses and keeps a tally of how many pulses have been seen until now, modulo 2*^N^* ; for example, when *N* =1 the counter simply computes the parity (odd or even) of the number of pulses seen so far.

The study of combinations of circuits and memory in living cells was initiated by Subsoontorn & Endy (2012<u>a</u>), Kim, Bojar & Fussenegger (2019), Siuti, Yazbek & Lu (2013), and Purcell & Lu (2014). A key step toward building artificial genetic memory was the design of the synthetic toggle switch (Gardner, Cantor & Collins 2000). Realizing the counting capability requires more elaborate logical operations. An early synthetic genetic counter that can count up to three induction events was based on transcriptional and recombinase-based cascades (Friedland, Lu, Wang, Shi, Church & Collins 2009). Also based on recombinases and information stored in DNA structure are the counters proposed by Subsoontorn & Endy (2012<u>b</u>) and more recently by Zhao, Pokhilko, Ebenhöh, Rosser & Colloms (2019). A pulse detecting circuit that responds only at the falling edge of a pulse was proposed by Noman, Inniss, Iba & Way (2016).

In this work, we present an *in silico* scalable and distributed counter design based on standard gene repressor networks, The design that we propose can be theoretically used to count modulo 2*^N^* for an arbitrary fixed integer *N* . However, practically implementing such an approach, even for very small *N*, faces a hurdle due to the practical impossibility of placing a large number of gates in single cells. It is known that large circuits can place stress on cells and overload them, causing failure of the design (Borkowski, Ceroni, Stan & Ellis 2016). This has led to the theoretical “retroactivity”, and to the construction of feedback mechanism to avoid their effects (Del Vecchio, Ninfa & Sontag 2008, Del Vecchio, Qian, Murray & Sontag 2018<u>a</u>). To avoid this issue, we base our design on distributed computation, with specialized cell types allocated to specific tasks, such as the computation of carry bits. This is an approach that we have also proposed for designing large classes of Boolean functions (Al-Radhawi, Tran, Ernst, Chen, Voigt & Sontag 2020). The communication between the different types of cells can be in principle implemented by diffusible small molecules such as quorum-sensing molecules. Distributed computation allows one to bypass constraints on the maximal possible number of circuit genes per cell, thus avoiding toxicity and making the design more scalable. (We assume relatively fast diffusion in a well-mixed environment.) To respect current experimental constraints (Gander, Vrana, Voje, Carothers & Klavins 2017, Nielsen, Der, Shin, Vaidyanathan, Paralanov, Strychalski, Ross, Densmore & Voigt 2016<u>b</u>), the maximum number of logical gates per cell is set to be seven. Our design is based upon a systematic and scalable optimization framework which aims to pick gate parameters so as to provide a degree of robustness, and which is especially suited to distributed implementations.

### Outline of approach

Our design process will follow the paradigm adopted in [11], which is well suited to the design of transcriptional circuits. Promoters and their associated genes are traditionally specified as pairs, or larger groups, in which a single or tandem promoter lies in front of a coding region. In this approach a biological gate consists of a gene coding region, together with the promoter region of a target gene that is regulated by the promoter of the another gene. Mathematically, the inputs and the outputs of each gate are promoter levels rather than repressor or protein levels.

Viewing gates in such a manner makes it easier to obtain expressions for a multi-input gate response, each quantified by its relative promoter strength. A more complicated cellular network can be made up of interconnections of such gates, each using promoter strengths as inputs and a promoter strength as its output.

Using the proposed paradigm, we design first a single-bit counter, i.e. a parity checker. When exposed to a sequence of *M* pulses of an external input (which might be a chemical inducer or a physical signal such as light at a specific frequency), the network will produce a binary output to indicate if the number *M* of pulses seen so far is even or odd. The single-bit counter uses an SR latch for implementing the memory function (Andrews, Nielsen & Voigt 2018<u>a</u>). Our design is asynchronous, meaning that it will detect pulse signals that are not equally spaced, though with a minimal separation time. The pulses themselves will be subject to specifications of minimal and maximal width. The single-bit counter serves as the key component of an *N* -bit counter, in which a count modulo 2*^N^* of the observed number of pulses is stored and displayed. The *N* -bit counter, as illustrated in Figure 1, can be constructed using *N* single-bit counters together with additional gates that implement the “carry” operations.

**Figure 1:**
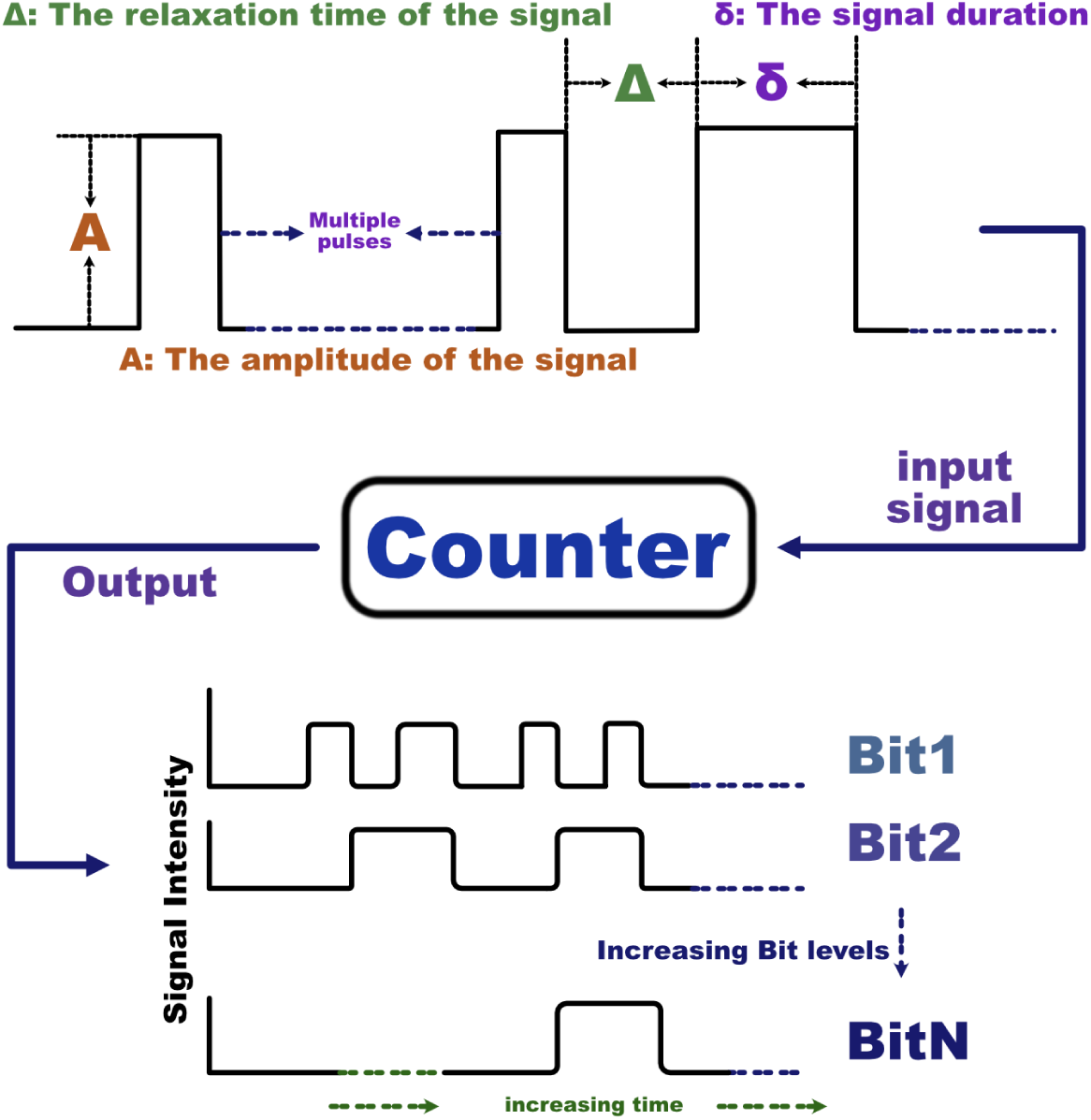
A schematic view of a multi-bit counter. A general *N* -bit counter takes an input pulse train as an input, and outputs a modulo-2*^N^* count of the total number of pulses, represented by the binary state of each bit (right). The pulse trains will be assumed to respect constraints on pulse width *δ*, spacing (relaxation time that permits the system to approach steady state until the next pulse.) between signals Δ, and pulse amplitudes *A*.

## 2 Results

### 2.1 The single bit counter circuit

#### Digital design construction

Using digital circuit design methods (Hardy & Steeb 2001, Jaeger & Blalock 1997, Bostock 1988), we propose a design for the single-bit counter as shown in Figure 2 (a).

**Figure 2:**
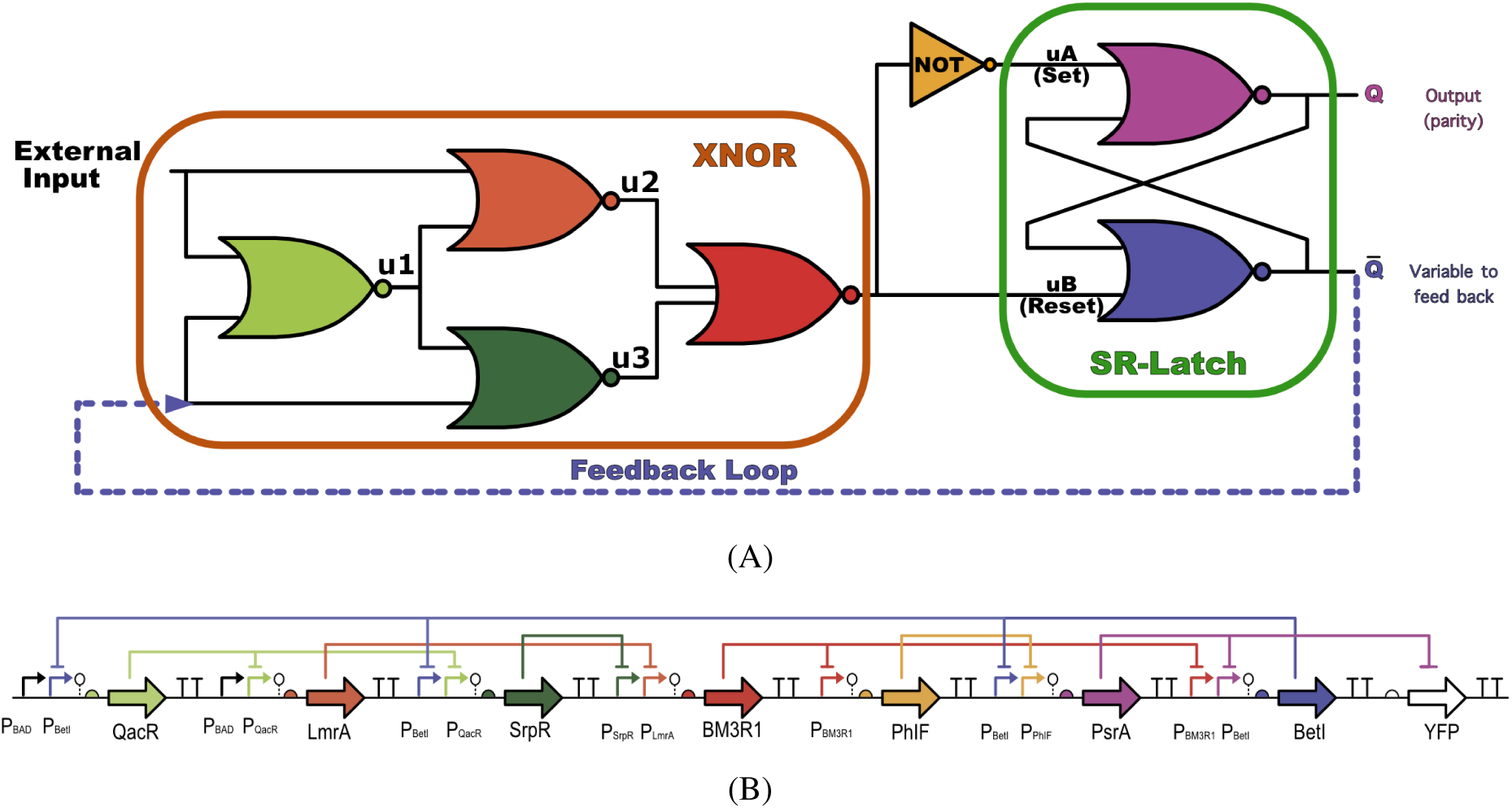
(A) Circuit design diagram for the single-bit counter. Shown is a seven NOR-gate implementation of the parity checker. This parity checker is a single-bit counter that contains two components. An XNOR component is shown inside the orange box, and an SR-latch component is shown in the green box. The design has a feedback loop from 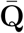 to the XNOR gate. The other input of the XNOR gate is the external input signal. The output of the single-bit counter is **Q**, which is the parity of the external input. (B) A genetic network implementation of a single-bit counter which consists of insulated gates. The diagram shows the promoter-gene-repressor network of one possible plasmid implementation in an *E. coli* system. Repressor colors shown in this genetic construct are matched with the gate colors shown in (A). All the terminators are shown as a black “TT”.

There are seven NOR gates in the design (the NOT gate is a NOR gate with the input repeated). This design consists of two components: an XNOR gate and an SR latch. One of the XNOR gate inputs is the external input to the counter, and in our application, it will consist of a sequence of pulses. The other input of the XNOR gate is fed back from one of the SR latch outputs. The SR latch module functions as a memory switch and the XNOR gate is a comparator whose output is one if and only if the inputs are identical.

The design in Figure 2 functions as a binary counter, in the following sense. Suppose that we start with the circuit at a steady state in which **Q** is low, and 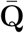 is high, which we think of as the “zero” or even parity state. This state should be stable. When an external signal changes from low to high, the output steady-state of the circuit will switch from (**Q**, 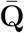)= (low,high) to (**Q**, 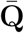)= (high,low), which is interpreted as “one” or “odd parity”. If the input signal goes back to zero soon enough, the circuit’s output will stay at this steady state, until the next input signal triggers a switch back to even parity, and so forth.

As an illustration, we show an example of how we would implement a single-bit counter in *E. coli*. By writing the binary state transition table for the logical circuit of the single-bit counter shown in Figure 2 (A), we have designed an *in silico* genetic network using Cello (Nielsen et al. 2016<u>b</u>, Andrews, Nielsen & Voigt 2018<u>b</u>) as an initial step. Then, we have used an optimization framework proposed in this paper to improve the aforementioned design by maximizing the range of admissible pulse widths. The final circuit is depicted in Figure 2 (B).

### 2.2 Circuit behavior

#### 2.2.1 Modeling framework

In order to simulate the behavior of the genetic circuit quantitatively, we write an Ordinary Differential Equation (ODE) model at the translational level. We use a standard Hill function to characterize the input-output response function of the gate (Nielsen et al. 2016<u>b</u>). For each gate, we use the following general ODE to model its dynamics:

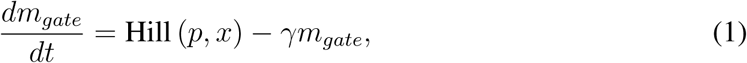

where *m_gate_, p, x, γ* are the concentration of the mRNA, the gate parameters, the inputs, and the mRNA decay rate, respectively. An archetypical Hill function and its parameters are shown in Figure 3.

**Figure 3:**
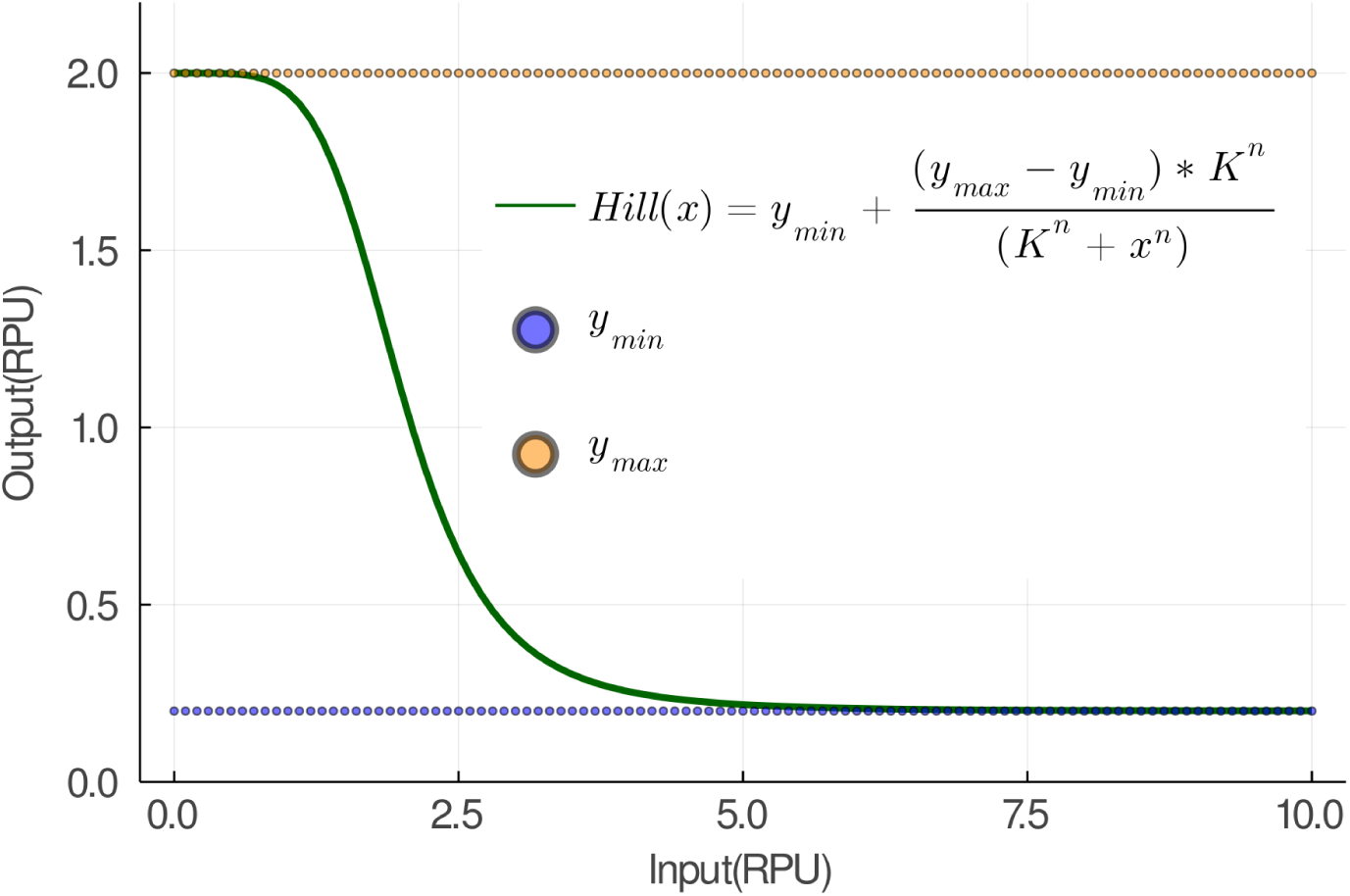
The response function of an arbitrary gate written as a shifted Hill function. The parameters *y_min_* and *y_max_* set the lower and upper bounds the response function output. The parameter *K* sets the width of the response range for the input. The parameter *n* controls the steepness of the response function.

The overall ODE model of the circuit is given by interconnecting the individuals gates via their inputs and outputs as is given the Methods section.

We checked through simulations that our design in Figure 2 (B) works as a parity-checker, see Figure 5. We study the stability of the single-bit counter and the circuit’s dynamical response to various external inputs signals in the following subsections.

**Figure 4:**
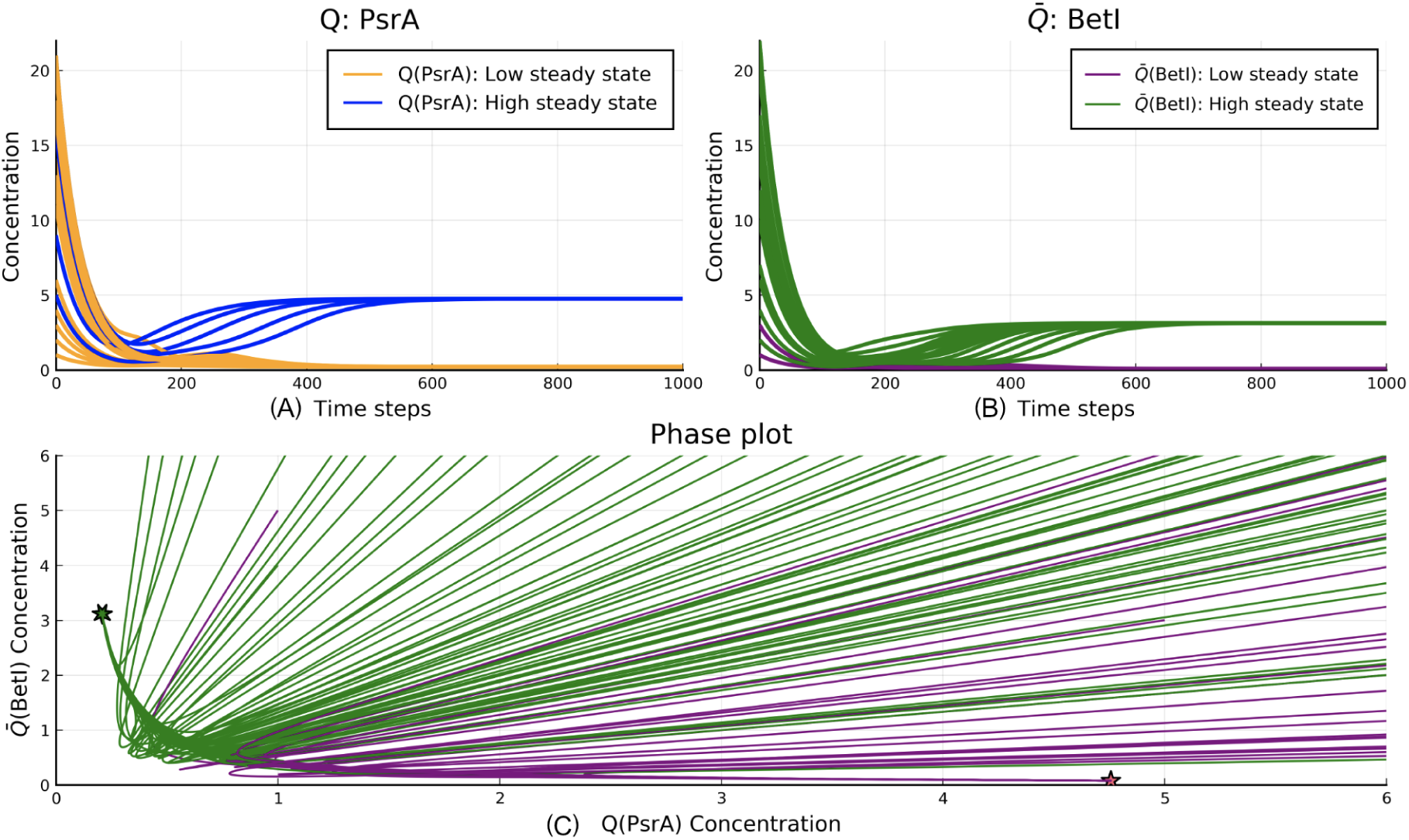
The bistability verification of the single-bit counter. (Panel A, B) With 50 randomized initial conditions, the SR latch states (**Q**, 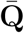, which determine the circuit’s output, asymptotically settle into one of the two steady states. The higher value steady state (for both Q or 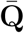) represents the binary ”1” state, and the lower value steady state represents the binary ”0” state. (Panel C) In the phase plot with random initial conditions, trajectories converge to one of the two stable steady states, represented here by two solid stars. Note that the intersecting trajectories represent projections of the original disjoint 7-dimensional trajectories. The two steady-state attractors are shown with stars in the (**Q**, 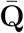) phase plane.

**Figure 5:**
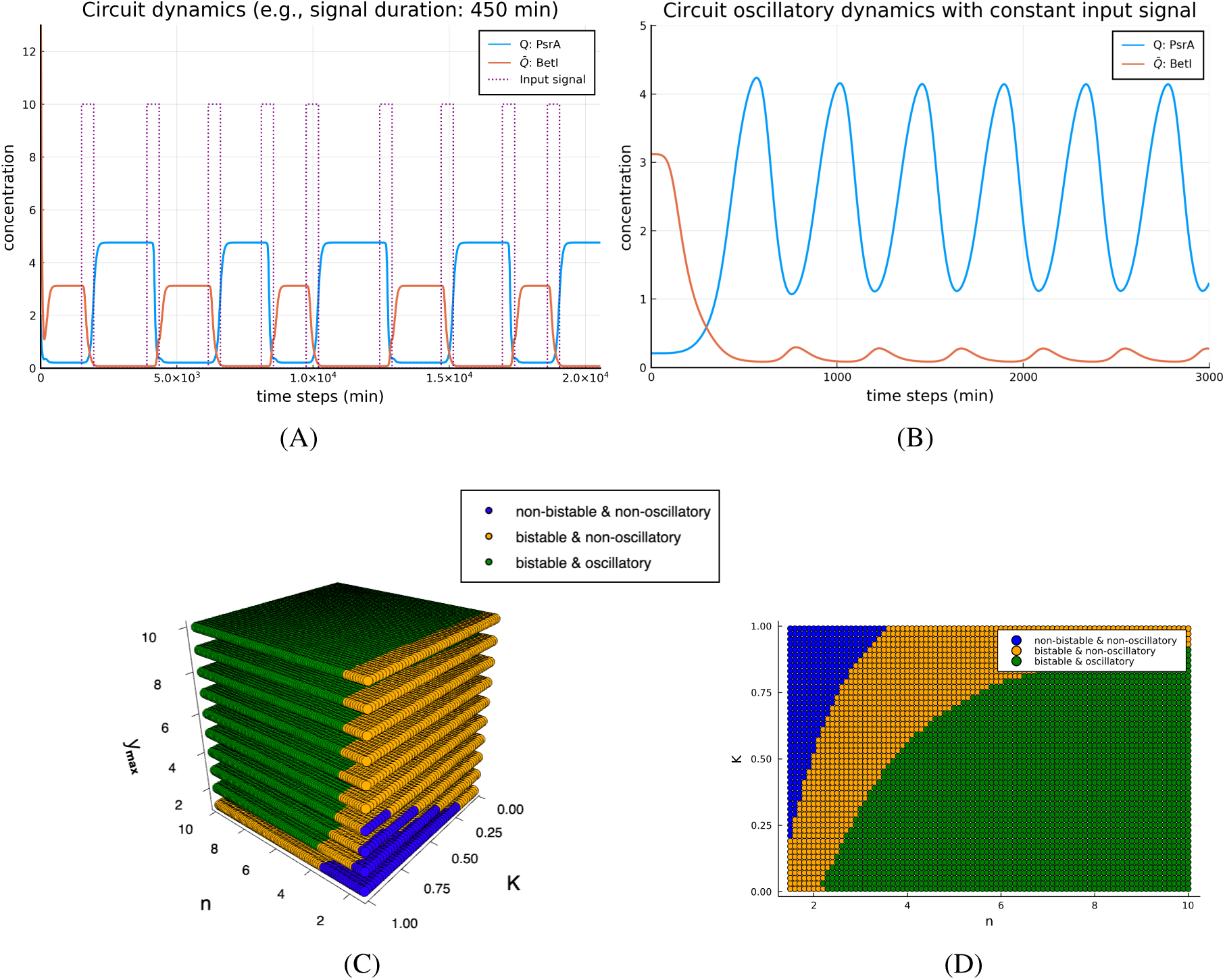
(A) The behavior of the single-bit counter under a pulse train input. The circuit’s output (represented by the gate **Q**) keeps track of the input’s parity correctly, as does the negated output 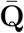. In this particular example, the input pulse has a duration of 450 minutes but with random relaxation times between adjacent pulses. (B) A demonstration of a stable oscillatory trajectory displayed by the counter when subjected to an external constant input. With the input’s amplitude *A* = 10, the circuit’s outputs **Q** and 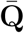 oscillate with 180*^◦^* phase difference. (C) A 3D bifurcation plot in the (*K, n, y*_max_) space. Each point is marked by whether the dynamics is bistable, oscillatory, both, or neither. (D) A layer of the 3D bifurcation plot in (C) with *y*_max_ = 1. *K*, *n*). Later in the paper, we will discuss designing the counter with gates that have different response functions.

#### 2.2.2 Circuit stability

The first property to be checked is that the system should reliably store the memory of the last state (even or odd) whenever the input signal stays at zero. This means that the autonomous (unexcited) system needs to be bistable, meaning that all solutions will generically converge to one of two states: high output or low output.

In the single-bit counter circuit shown in Figure 2 (B), a system of the ODEs with seven state variables describing a seven-repressors circuit is used for modeling the dynamics of the circuit evolution. For the output gates of the circuit, before understanding how its dynamics is affected by an external input signal, one needs to check first whether the outputs of the circuit produce binary values. In order to numerically check the bistability of the circuit, we randomly initialize all the seven state variables (repressor gate concentrations) and simulate the trajectory of a 7- dimensional differential equation until it settles on a steady state. The system’s outputs are the states of the SR latch: **Q**, 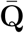. Figure 4 shows the projected trajectories of (**Q**, 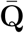) where all states asymptotically converge to either a high **Q** (PsrA) or a high 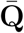 state (but not both). (BetI) value at steady

#### 2.2.3 Counting capability

A functioning single-bit counter must switch outputs if the input stays high for a sufficiently long duration of time. The duration of the input signal pulse *δ* and the relaxation time Δ (when the signal is off) are both critical factors that affect the correct functioning of the parity checker (Figure 1). It is necessary for the pulses to be long enough to allow the SR-latch to flip, yet not too long, since a long input will flip the latch back to its original state. Similarly, the interval between pulses should be large enough to allow the protein concentrations to settle into their steady states.

A key part of the design concerns the characterization of the types of pulses that the counter can reliably count. This means finding the ranges of the two parameters *δ*, Δ (Figure 1), which are the pulse duration and the relaxation time, respectively. To that end, we have performed simulations to estimate the empirical duration of the constant input signal that guarantees that when the input pulse turns from high to low, the system stays in the newly switched steady state. Using an optimization method, we found that when the pulse duration is in the range of 300 to 550 minutes, the switching behavior is persistent and stable. We will revisit the issue of parameter selection later in the paper.

Figure 5 (A) shows an example illustrating how our circuit responds to an input which is a square wave signal with a fixed duration and randomly varying relaxation times. When the circuit is at a steady state, and an input signal is applied, the system output switches state. When the input signal is at a low value, the circuit stays in the new switched state until a new pulse arrives to switch the output to the previous steady state.

##### Constant inputs lead to oscillations

In our proposed design, a high external input continuously tries to flip the output as per the state transition diagram (Methods, Table 1). Hence, we expect the output to show a stable oscillatory behavior when the external constant input stays high. Such a design is reminiscent of the standard oscillator architecture consisting of a negative-loop with a delay. We indeed confirmed the oscillatory behavior of the circuit by simulations in Figure 5 (B).

**Table 1:** State transition table of the open-loop circuit.

| X | Y | Q | $\bar{Q}$ |
| --- | --- | --- | --- |
| 0 | 0 | 1 | 0 |
| 0 | 1 | 0 | 1 |
| 0 | 0 | 1 | 0 |
| 0 | 1 | 0 | 1 |
| 1 | 1 | 1 | 0 |
| 1 | 0 | 0 | 1 |
| 1 | 1 | 1 | 0 |
| 0 | 1 | 0 | 1 |
| 0 | 0 | 1 | 0 |
| 1 | 0 | 0 | 1 |

Furthermore, we have extensively explored the oscillatory behavior of the circuit dynamics for a constant input signal. The kinetic constant *K*, the cooperativity index *n*, and the maximum value *y*_max_ of the Hill function are the three most important parameters for the circuit dynamics. A 3D bifurcation plot in the phase space spanned by *K*, *n*, and *y*_max_ values is demonstrated in the Figure 6 (C). Our simulations show that the phase space has three distinct regions, which are “no oscillation and no bistablity”, “no oscillation and bistablity”, and “oscillation and bistablity”.

**Figure 6:**
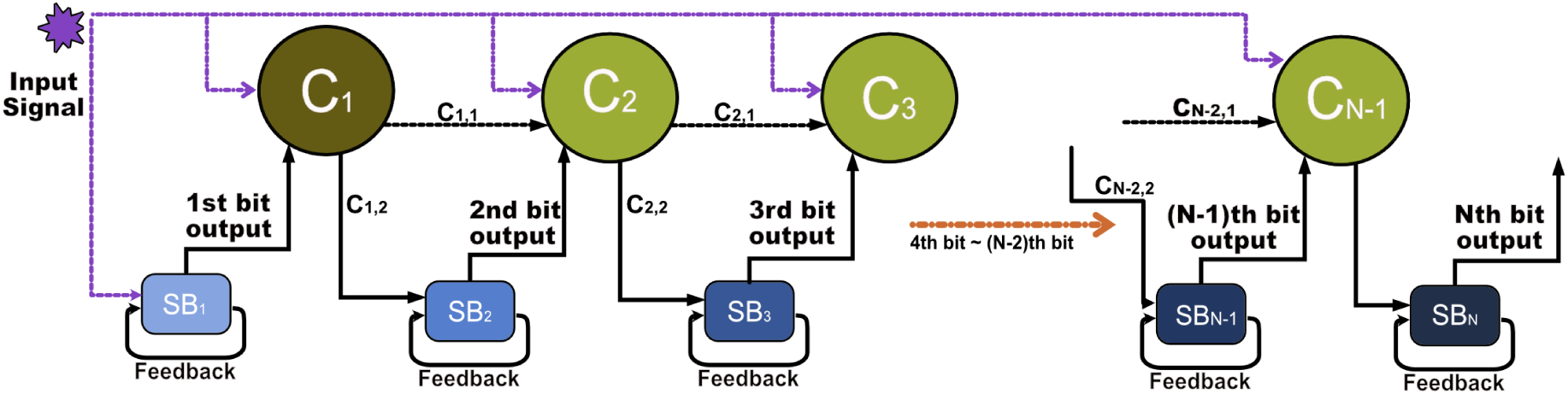
Schematic design of a generic modular multi-bit counter. The counter employs three types of modules, each implemented in a different cell type. One connector of type 1 and *N* 2 connectors of type 2 (colored in dark and light green respectively) are shown in the first row and *N* single-bit counters (colored for different bit levels gradient blue) are in the second row. The external input signal is colored in dashed purple lines, and it acts on all connectors simultaneously. Black arrows denote communication via diffusible small molecules. Each connector outputs two signals *C_i,_*_1_, *C_i,_*_2_ at *i*th bit level. The *C_i,_*_1_ is the AND operation result between the previous bit’s output and the previous connector’s output. The *C_i,_*_2_ is the AND operation result between the *C_i,_*_1_ and the external signal.

### 2.3 Multi-bit counter

In the previous section, we have proposed a genetic circuit design for a single-bit counter. This design functions as an universal parity checker that can be re-used in a multi-bit counter design.

#### 2.3.1 Design principle

We design the multi-bit counter system in a highly modular fashion. Using a single-bit counter as the building block module, we construct a multi-bit counter by adopting the following pattern: **1-bit counter – Connector – 1-bit counter**. Hence, a single-bit counter will be implemented repetitively as a module, which simplifies the design process. We will also design connectors to interface single-bit counters for intercellular communication. The overall design can be scaled up, in principle, to any arbitrary number of bits by interconnecting single-bit modules and connectors modules. Such a scaled-up design can be achieved as long as enough orthogonal diffusible small molecules such as quorum-sensing molecules are available.

Our design of the multi-bit counter enjoys a degree of robustness, which we will quantify using the trade-off between the mean value of the pulse duration time (*δ*-mean) and the standard deviation of the set of all possible pulse duration times (*δ*-std). In order to avoid a computationally-infeasible combinatorial search among different gate parameters, we first mathematically study the idealized case in which every gate has the same characteristics (Hill coefficient, *y_min_*, *y_max_*,

#### 2.3.2 Modular design construction

Following the general principle outlined above, we propose the following scheme for constructing a multi-bit counter system. Any design of the multi-bit counter system consists of the following three essential modules. Different modules talk with each other via diffusible small molecules.

a. **Module 1: The parity checker**. This module is the building block for constructing a multi-bit counter system, and it is responsible for storing and updating the value of the corresponding bit. For example, if one wants to build a 3-bit counter, we will use three copies of the first module, and each one will be used for one of the bits.
b. **Module 2: Connector type 1**. This specific module uses one diffusible small molecule connecting the first parity checker and the second parity checker only. As shown in Figure 6, the inputs are the external signal and the 1st parity checker’s output. The output of the connector will be communicated to the parity checker and the connector at the next stage by diffusible small molecules.
c. **Module 3: Connector type 2**. This module serves as the universal connector that communicates between the remaining parity checkers beyond the 2nd parity checker.

With the proposed three modules, Figure 6 shows a schematic view of the design of a generic *N* -bit counter.

Given the proposed design scheme, the next step is to find the Boolean representation of the two types of connectors. Figure 7 shows how to find the corresponding Boolean function for connectors using a truth table. We illustrate this process by finding the connector type 1 Boolean function as an example. The figure lists all four stages of the possible states of a 2-bit counter and the state transition table associated with each stage. For example at stage 2 in Figure 7, the counter is at its internal state *B*_2_*B*_1_ = “0 - 1”. When the input signal changes from “0” to “1”, the 2-bit counter should switch from “0 - 1” to “1 - 0”. The connector takes the external signal and the first bit as inputs, and its output will be “1” when its inputs are “1- 1”. Similarly, we can repeat the same process for the rest of the counting situations to derive the desired output for all possible inputs. As for the simplest case, we can directly see that the Boolean function of the connector type 1 is just an “AND” gate. Similarly, the Boolean function that represents the connector type 2 can also be determined from the requirement of the binary representation of the state transition table at the higher connector levels beyond the 2nd-bit level.

**Figure 7:**
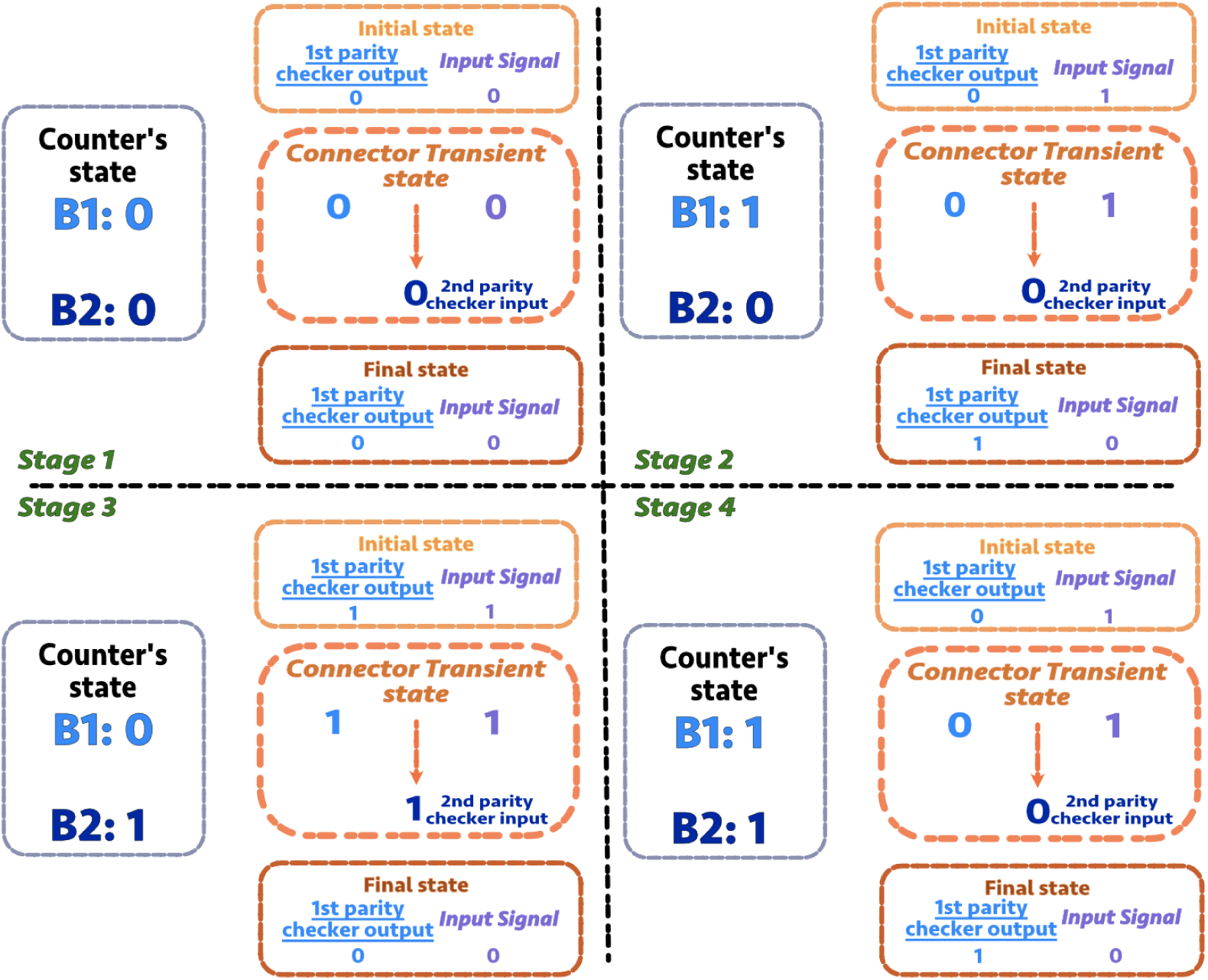
Connector type 1 state transition table. Four stages of the 2-bit counter are shown. The state of 2-bit counter can be represented by the expression of **B1** 2^0^+**B2** 2^1^. **B1** represents the 1st-bit binary state, and **B2** represents the 2nd-bit binary state. **B1** and **B2** are either “1” or “0”. On the right side of each quadrant in the orange outline, the connector transient state table shows how the 1st parity checker’s output and the external signal input change the 2nd parity checker’s input. Each stage (each quadrant in the figure) contains four pieces of information: 1) The binary representation of the steady state for each bit (the 1st-bit is denoted by **B1** and the 2nd-bit is denoted by **B2**). 2) The initial state of the 1st-bit output and the input signal. 3) The connector transient state (the middle orange dashed circle). 4) The final state of the 1st-bit output and the input signal.

With the connector type 1 functioning as an ”AND” gate, one is able to implement such a connector with two NOT gates and one NOR gate as shown in Figure 8 (A).

**Figure 8:**
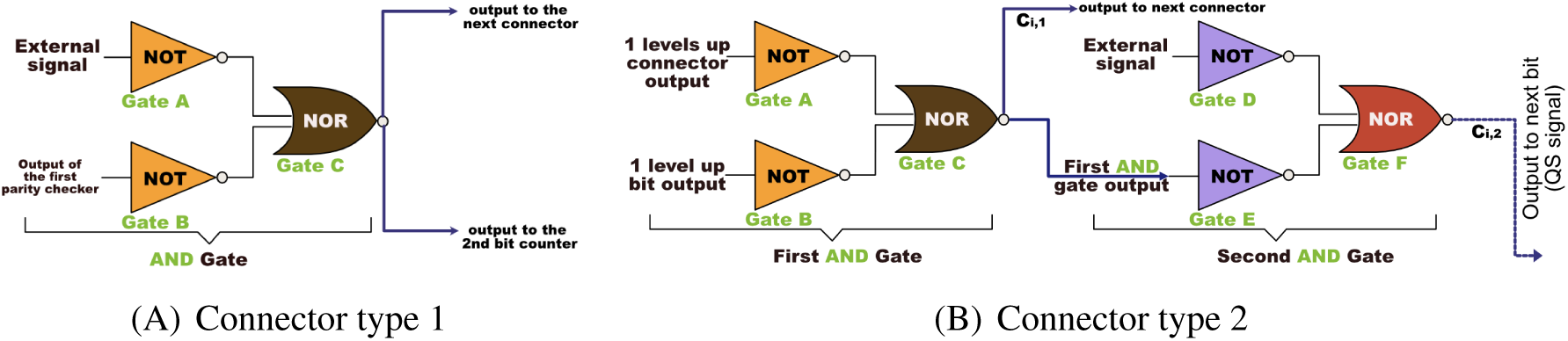
(A) Connector type 1. This module will be used to interface the first and the second parity checker. The depicted design implements an AND gate using a 2-input NOR gate. (B) The connector type 2 module design. The Boolean expression for this module in the multi-bit counter design is represented by a cascade of two AND gates. The first AND gate integrates the signals from the outputs of the preceding two parity checkers. The output of the first AND gate will be one of the inputs of the second AND gate. The other input for the second AND gate is the external signal. The connector 2 output will use a diffusible small molecule for inter-cellular communication.

#### 2.3.3 Schematics of the 2-bit counter

In Figure 9 (A), we show the schematic design for a 2-bit counter that is constructed by combining two parity checkers and a connector type 1. Figure 9 (B) demonstrates the counting capability of a design for pulses that fall within the proper ranges.

**Figure 9:**
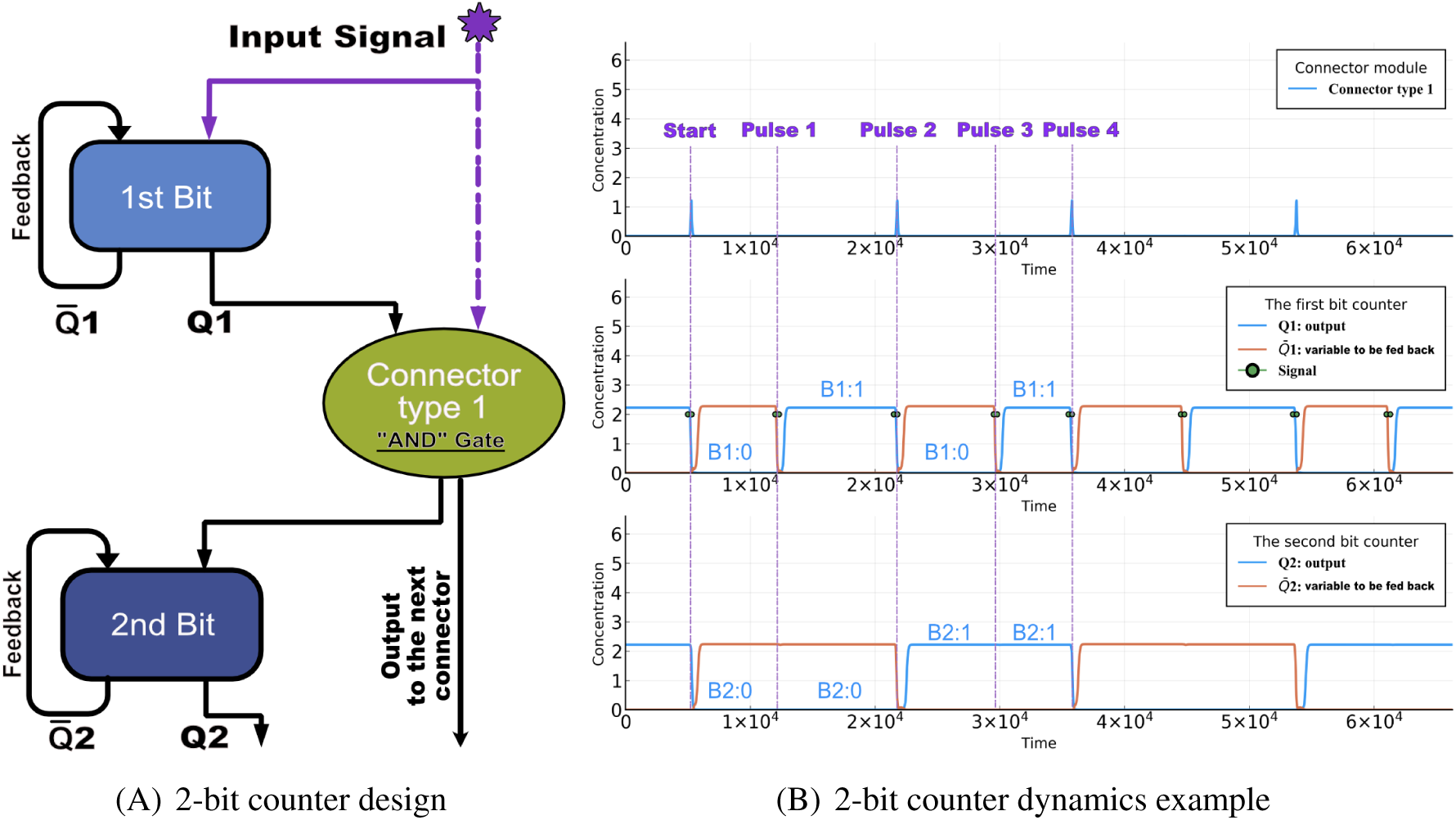
The 2-bit counter design. (A) Schematic design of the 2-bit counter. The signal **Q**1 is the output while 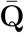1 is the feedback signal for the 1st parity checker. The connector type 1 takes **Q**1 and the external input signal as inputs. The output of the connector module can be a diffusible small molecule which serves as the input for the 2nd parity checker. (B) We show an example trajectory of each module aligned with the schematic design in (A). The counter starting time is determined by the first time that the last bit switches state, as explained in the text.. Due to the plotted long relaxation time (relative to input pulse width) between adjacent pulses, each green dot in the graph, if zoomed in, represents one input pulse as shown in Figure 5. The representation of the 2-bit counter ”**B2**-**B1**” is used to indicate the output state “**Q**2-**Q**1”

The persistent comb-like structure in the first row indicates that the carry bit information of the 1st-bit has been successfully transmitted to the 2nd-bit. To examine if the chosen parameter set can give rise to a working 2-bit counter, we first randomize the initial conditions for all state variables in the 2-bit counter dynamical equations and wait for the counter to equilibrate to one of its steady states. Then we give the system an external input signal, which is a train of short pulses. To capture the nature of the input signal’s noisy environment, we add noise to the pulse widths to simulate asynchronous counting scenarios, which would mimic more realistic biological situations. Specifically, we allow the Δs to be randomly sampled from the range of [Δ_0_, 2Δ_0_], with a given relaxation time Δ_0_.

A 2-bit counter will follow the pattern of “0 - 0”, “0 - 1”, “1 - 0”, “1 - 1”, and then return to “0 - 0”, which corresponds to counting input pulses from 0 to 4. As illustrated in Figure 9-b), we show a full cycle trajectory of a 2-bit counter. Each pulse that presents as an input to the counter will increment the internal state of the counter.

#### 2.3.4 Initialization Scheme

To determine the initialization scheme for a multi-bit counter, we consider two issues. The first one is whether the counter has two distinct steady state solutions for the output Q. As demonstrated in Figure 5-C,D, the circuit is bistable for a wide range of parameters, and all the trajectories (with randomly-chosen initial conditions) converge to a steady state. The second issue is when should counting start and what constitutes a zero. This is a major issue when designing dynamical circuits. The problem is that it may be difficult to set a system, and our counter in particular, to a desired initial state. We have solved this problem as follows. The system should be preconditioned by giving it a sequence of training pulses. The initial time is then set as the first time that the last bit switches. Extensive simulations with random initial conditions show that this always results in the correct initialization after a few (no more than 5 for most of the time, for a three bit counter discussed below) pulses. An alternative approach is to define “zero” as the initial state of the counter, and the actual events counted can be computed as the difference between the state of the counter at the current and the initial state. Yet another approach would be to include a “reset button” in the design, at the cost of adding a constitutive promoter that is controlled by an additional inducer which, when applied, sets the initial state.

#### 2.3.5 Schematics of the 3-bit counter

We take the previous design a step further and showcase a 3-bit counter built by implementing an additional connector type 2 and a 3rd single-bit counter on top of the 2-bit counter system.

We introduce the connector type 2 as a universal connector module for connecting single-bit counters beyond the second single-bit counter in a general multi-bit setup. Figure 8 (b) shows a schematic view of the connector type 2 using NOT and NOR gates. Similar to the connector type 1 module, we can identify the Boolean function that the connector type 2 represents. In a 3-bit counter system, the corresponding transient state table for the Boolean function is similar to the one depicted in Figure 7, but is much larger as it consists of 8 stages. The result is that a connector type 2 consists of a cascade of two AND gates. Each AND gate is realized by 2 NOT gates and 1 NOR gate. The first AND gate takes two inputs, which are the outputs of the two preceding parity checkers. The second AND gate takes the output from the first AND for the counter. The parameters for this working 2-bit counter are: the scaling factor *ξ* = 0.025, the degradation rate *γ* = 0.025, the equilibrium constant *K* = 0.011, the cooperativity index *n* = 1.5, the duration of external signal *δ* = 300, the amplitude of the external signal *A* = 20, the shifted hill coefficient maximum value *y_max_* = 3, the shifted hill coefficient minimum value *y_min_* = 0.002, and the initial and between signals relaxation times Δ_0_ = Δ = 5000. gate and the external signal as inputs and produces the connector type 2’s output for the next bit level by using a diffusible small molecule.

Similar to the 2-bit counter, we show a schematic design and the dynamics for each module of the 3-bit counter in Figure 10. An example of a cycle that successfully counts the number of pulses from 0 to 8 is shown in Figure 10-B). As with the two-bit counter, the initial time is set by the first time that the last bit switches, after a preconditioning set of pulses. Therefore, the initial counter state “0-0-0” is defined as the state during the times between the dashed lines labeled “start” and “P1” (see Figure 10-B).

**Figure 10:**
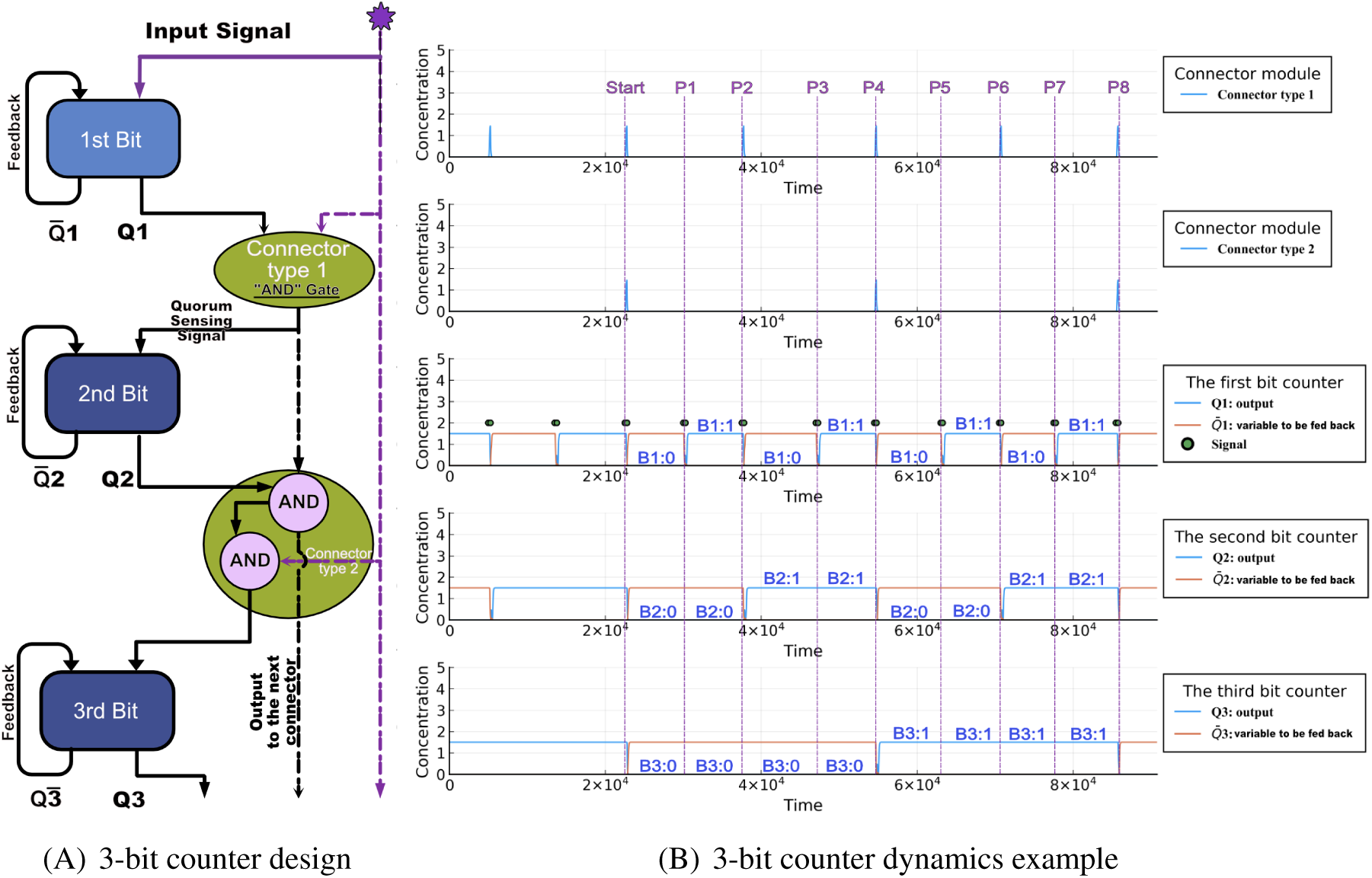
The 3-bit counter design schematics. The inputs are shown in the middle panel, for ease of reference. (A) The diagram shows a modular design of the 3-bit counter following the pattern of “counter-connector-counter”. The only difference between the 3-bit counter and the 2- bit counter is that the second carry bit module uses the connector type 2. **Q**1, **Q**2, and **Q**3 are the outputs of each bit level, and 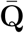1, 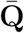2, and 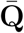3 are the variables to be fed back. (B) We show an example trajectory for each module. The starting time of the 3-bit counter is set by by the first time that the last bit switches state, as explained in the text. The steady state binary representation for each parity checker **B1**, **B2**, and **B3** are marked on top of the gates’ trajectories. The parameters for this 3-bit counter example are: the scaling factor *ξ* = 0.025, the degradation rate *γ* = 0.025, the equilibrium constant *K* = 0.081, the cooperativity index *n* = 2.81, the duration of external signal *δ* = 270, the amplitude of the external signal *A* = 20, the shifted hill coefficient maximum value *y_max_* = 1.5, the shifted hill coefficient minimum value *y_min_* = 0.002, and the initial and between signals relaxation times Δ_0_ = Δ = 20000.

For a multi-bit counter system with more than 3 bits, one can easily generalize the counter system and build upon the 3-bit counter by adding additional modules of connector type 2 and single-bit counters as we have demonstrated in Figure 6.

##### Scalability of the counter

So far, we have designed a single bit counter, a 2-bit counter, and a 3-bit counter. The design principle that we proposed can be applied to the design of a distributed *N* -bit counter. As a demonstration, we showcase the dynamics of an 8-bit counter in Supplementary Figure 5.

For a large *N* -bit counter, the variation in the timing of the input signal that reaches to each module (a connector or a parity-checker) can be an important factor when building a feasible counter experimentally. Depending on whether the actual cellular colonies are grown on agar or in a mixed liquid culture, the tolerance of the variation for the input signal for each bit level may depend on the spatial distribution of each module’s cellular colony, but also on the viscosity of the liquid, the temperature, and the Hill coefficients of the gates.

### 2.4 Design selection graphs

In the previous sections, we described a generic way of constructing a multi-bit counter. We have simulated wide ranges of parameter values and produced databases for different multi-bit counters (2-bit and 3-bit counters), see Figure 11. In the generated database, many parameter sets can give rise to a working counter. However, in synthetic biology, experimentalists often have a limited selection of genetic gates, and these gates usually respond to a restrictive dynamic range of input. For simplicity, we first assume all the activation functions to be identical. Then, we sample gates from the generated multi-bit counters’ database to produce generically working counters in which each gate has its own activation function.

**Figure 11:**
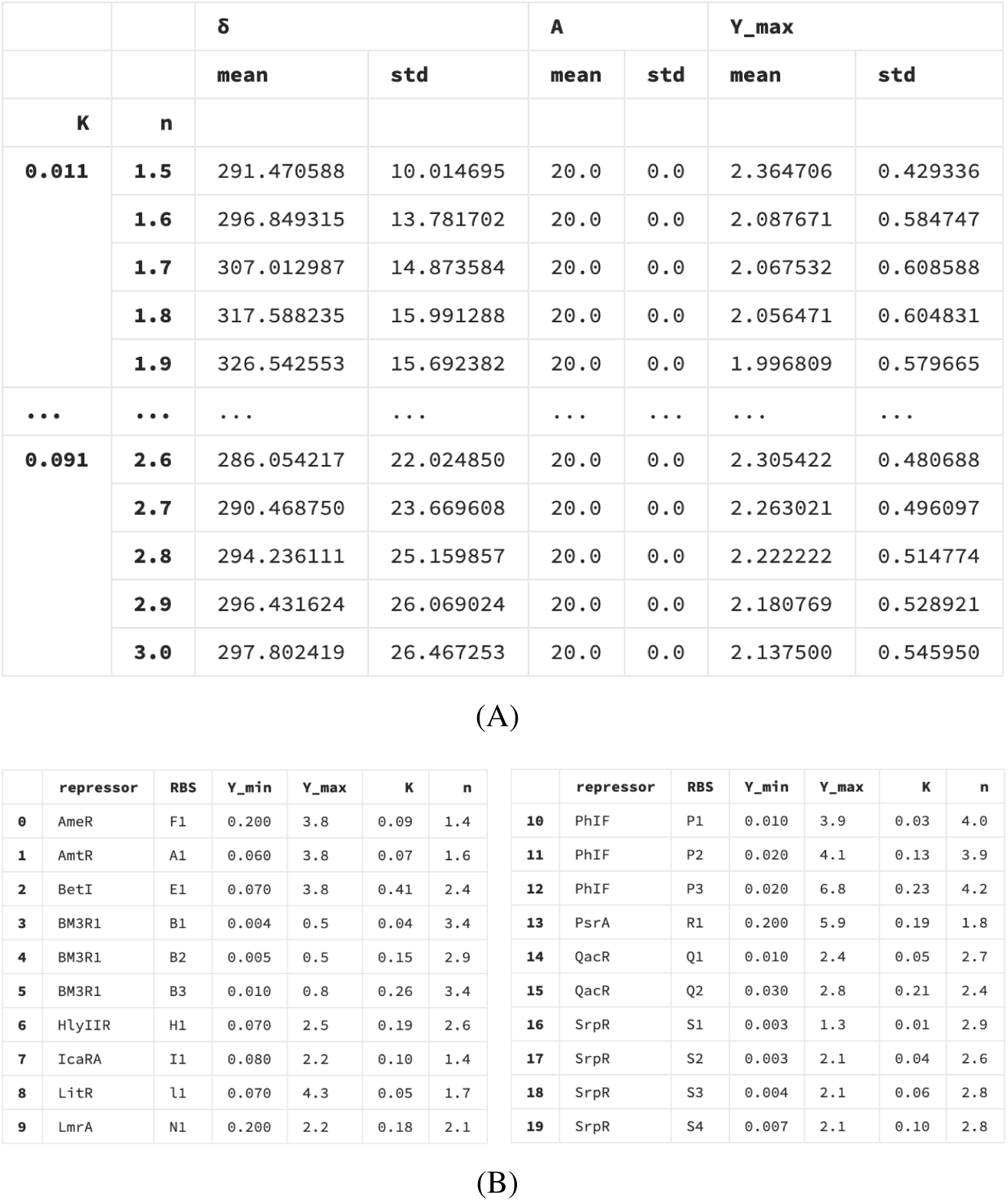
(A) The optimization results for the 2-bit counter. For each pair of (*n, K*), the table hierarchically displays the values of the mean and the standard deviation of the pulse widths *δ*s, and the *y_max_* value of the Hill coefficient for a given external pulse train with amplitude *A* = 20. Due to the simulation time constraint, we only simulated discrete values of *K* and *n* with increment by 0.01, and 0.1 respectively. The “…” line represents the part of the database which is not displayed here. The complete table can be found in the github link https://github.com/danielchen26/circuit-design. (B) A library of parameters table for a list of synthetic transcriptional gates in *E. coli* (Nielsen et al. 2016<u>b</u>) .

From an experimental point of view, it is desirable to pick genetic components so as to obtain a robust counter. Towards this goal, we propose a method for optimizing the selection process when limited genetic components are available. Such a process will allow designing the counters based on availability of parameters. The parameters that are essential for constructing a feasible multi-bit counter are the shifted Hill coefficients *y_min_*, *y_max_*, *K*, *n*, the signal amplitude *A*, and the pulse width *δ*. A general response function (shifted Hill function) has been illustrated in Figure 3.

We make design selection graphs that can help with experimental design in two ways. First, given a physical genetic gate with a kinetic parameter (*K*) and a cooperativity index (*n*), our formalism will estimate an optimal operating range for the input signal and how robust the pulse width will be against perturbations. Second, if we instead are given desired durations of external signals, we can find a range of parameters of a genetic gate that results in a functioning counter. If such a gate is not yet available, this will suggest how to design a new synthetic gate according to the desired Hill parameter sets.

For each parameter set of the Hill function, we sample many possible input signals with different amplitudes and durations. We then calculate the mean values and the standard deviations of the subset of the simulated pulse widths (*δ*) that gives rise to a working counter as shown in Figure 11 (A). Figure 11 (B) shows currently available synthetic gates in the *E. coli* system, and each gate is associated with a unique (*n, K*) pair. For instance, given the current knowledge of the available gates listed in Figure 11 (B), one can select an optimal gate parameter set given constraints that *n <* 3, *K <* 0.5 that has the highest robustness. By “robustnes,” we mean: given a mean value of a pulse width *δ*, the larger standard deviation around this mean value, the more robust the counter would be, because the larger value of *δ* standard deviation corresponds to more available counters in the (*n, K*) space. Given a particular set of parameters for the gates, one can also obtain the optimal pulse width for assessing the counter’s robustness.

Once given available genetic gate characteristics, represented by the (*n, K*) pairs, one can refer to Figure 11 to find the mean external pulse width and the mean level of *y_max_* that is required for a design. To quantify how robust a given genetic gate is, the standard deviation of *δ* and *y_max_* can provide useful information about the tolerable variations in the duration of the external signal and in the maximum value of the Hill activation function.

In order to guide gate design, we present the data alternatively in Figure 12 for the 1-bit, the 2-bit counter, and the 3-bit counter. It can be noticed that the mean of the pulse width in each counter changes wildly in the (*n, K*) plane. On the other hand, when comparing the overall shift in the whole (*n, K*) space, the signal mean distributions are not dramatically different between the 1-bit and 2-bit counters. For those synthetic gates with low cooperativity indices, the counter will operate with signals with lower values for the mean of *δ*. As for the trend of the standard deviation of *δ*, we can see from the 2nd column that robustness can be gained by increasing the cooperativity index *n*.

**Figure 12:**
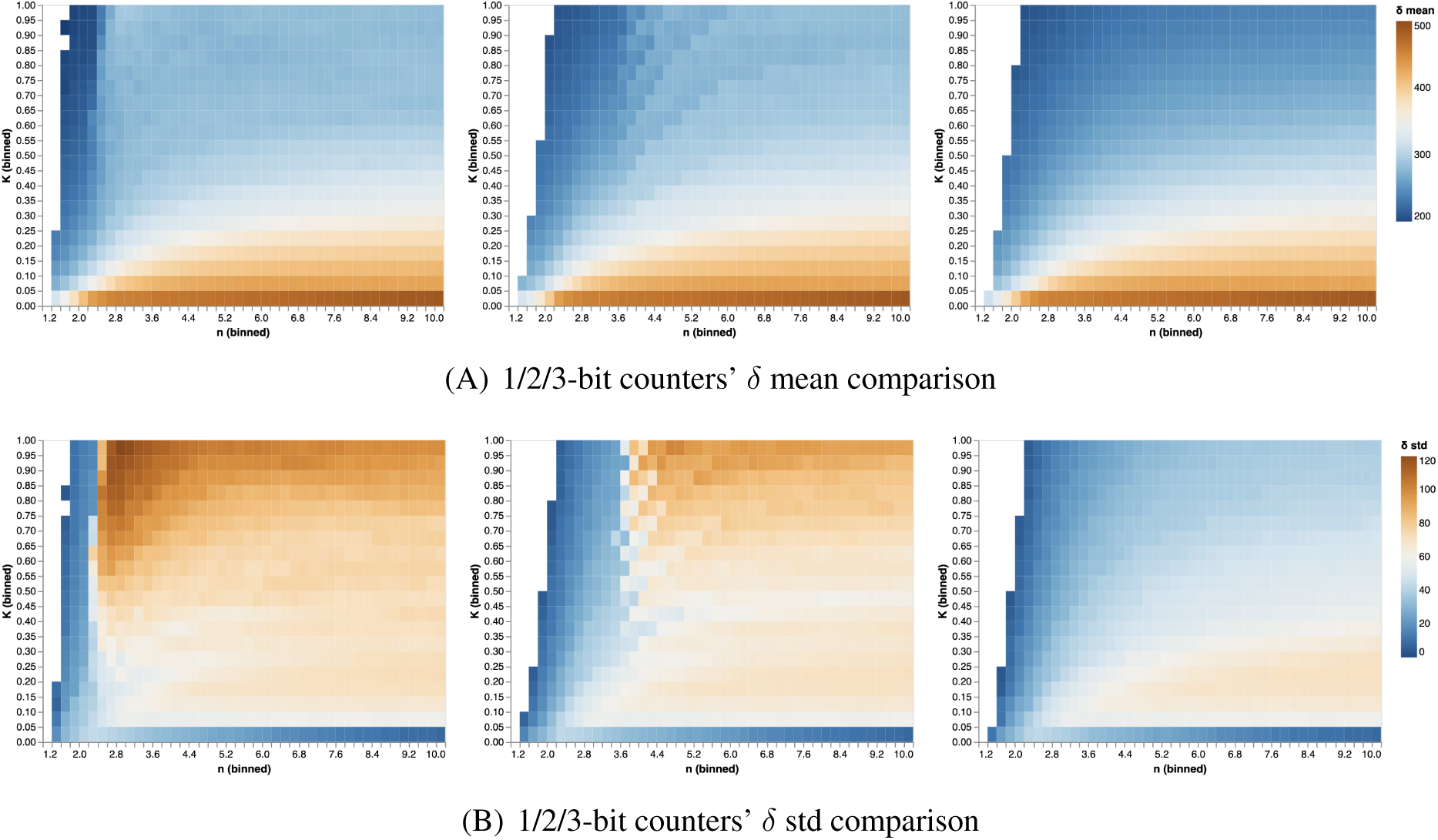
The mean and the standard deviation of the pulse width (*δ*) for the 1-bit counter, the 2-bit counter, and the 3-bit counter for various (*n, K*) pairs. We explore the (*n, K*) space with the cooperativity index *n* ranging from 1 to 10, and the kinetic constant *K* ranging from 0 to 1. The pulse width’s mean (*δ*-mean) for each (*n, K*) pair indicates the mean of the feasible range of pulse widths for a specific (*n, K*) pair. The pulse width’s standard deviation (*δ*-std) quantifies the robustness of a given counter against perturbations. The plots show that the 1-bit counter can operate with smaller values of *n*. The region of robustness for the 2-bit and the 3-bit counters shifts rightward in the (*n, K*) plan compared to the 1-bit counter.

In order to understand the trade-off between the mean pulse width (*δ* mean) and gates’ robustness and to understand how such a trade-off evolves as we scale up to a multi-bit counter situation, we show a comparison graph for the 1-bit and 2-bit counters in Figure 13 as an illustration. In this graph, we plot *δ*-mean vs. *δ*-std for the 1-bit, the 2-bit counter, and the 3-bit counter, and show the trend of how they shift in the (*n, K*) plane.

**Figure 13:**
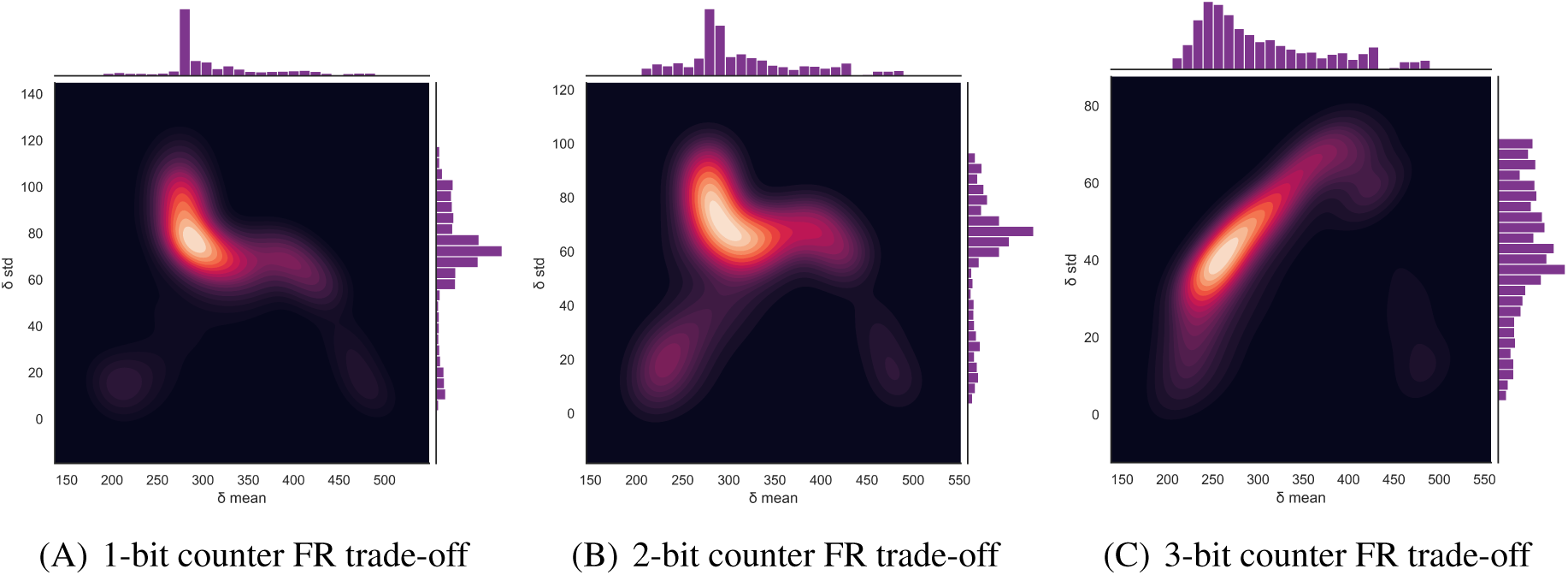
Feasibility vs. robustness (FR) trade-off comparison between the 1-bit counter, the 2-bit counter, and the 3-bit counter. For all simulated synthetic gates, we plot a 2d density graph with contours for *δ* mean vs. *δ*-std. The plots help identify the preferred region for designing new synthetic gates. The contour lines represent different levels of probability density for data points. The darker, the higher probability it represents. For the single-bit counter, the best region for designing a synthetic gate is around *δ*-mean = 300 and *δ*-std = 75. For the 2-bit counter, the best region for designing a synthetic gate is *δ*-mean from around 260 to 350, and *δ*-std can range from 60 to 90. For the 3-bit counter, the best region for designing a synthetic gate is around 240 to 320, and *δ*-std can range from 20 to 70.

To see how our design selection graphs shed light on experimental synthetic gate design, we compare the available experimental gate parameters with the above analysis. To gain insight for the two commonly used systems, Figure 14 shows the transcriptional gates available for both the *E. coli* and yeast systems in the simulated (*n, K*) space as shown in Figure 12.

**Figure 14:**
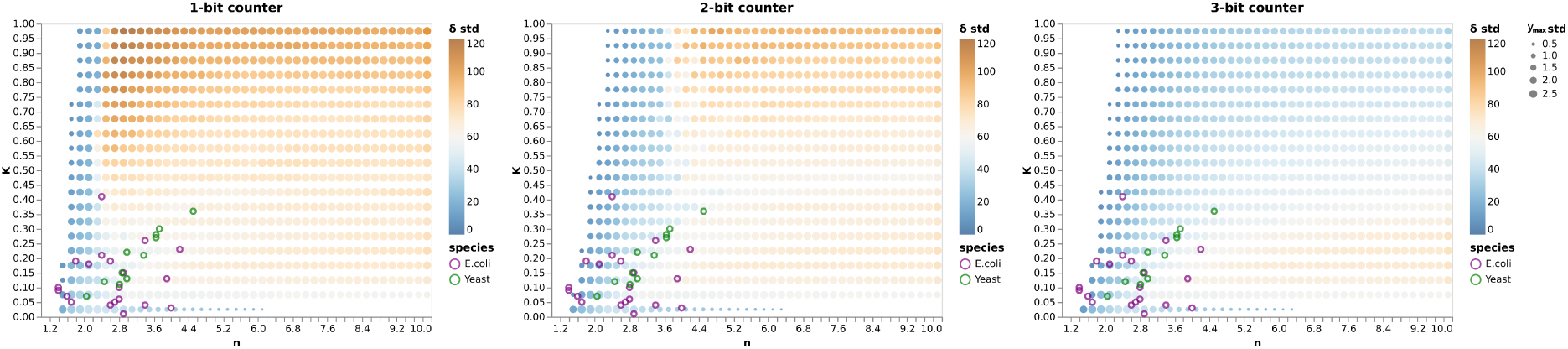
Experimental transcriptional gates in the simulated (*n, K*) space. We plot experimentally built synthetic gates for the *E.coli* and yeast system on top of the *δ*-std plane for both 1-bit counter, 2-bit counter, and 3-bit counter as an example. The purple circles represent the experimental values of the synthetic gates for the *E. coli* system, and the green circles are for the yeast system. The *δ*-std is colored with the solid dot, and the size of the dot represents the standard deviation of the *y_max_* value. We have sampled *n* from 1 to 10 with a 0.2 step size, and *K* from 0 to 1 with a 0.05 step size in the simulations.

Figure 14 shows that most synthetic gates built for the *E. coli* system display, on average, less robustness than the synthetic gates built for the yeast system. The most robust gate of all experimental gates seems to be the yeast gate with *n* = 4.6 and *K* = 0.35. The plots also show that the currently-available physical gates are not in the most robust region of the space, which leaves lots of room for improvement in synthetic gate design.

From the perspective of designing synthetic gates, the above comparison between the experimentally used genetic gates and the more robust simulated gates can provide guidance when designing new synthetic transcriptional gates.

#### Different activation functions

The above study covered the ideal case in which the counter system can be built out of gates with identical activation functions. However, typically one must use gates with different parameters. Next, we show a generalization of the aforementioned design with the 1-bit counter database as an example, which allows for implementing gates with different activation functions.

In Figure 15, we show an example of how one could easily generate various instances of a 1-bit counter in which each gate is associated with a different activation function, specified by (*n, K*). Given a pulse width *δ*, we search our generated databases for seven gates that are *simultaneously feasible*. This means that the feasible ranges of pulse widthes of the seven selected gate have a maximal intersection. We can see that as long as the seven selected gates have an overlap in their *δ* distributions, then a feasible 1-bit counter can be realized using the selected parameters. For example, we found that, in general, if one only focuses on the range of *δ* from the 25% quantile to 75% quantile for each gate’s *δ* distribution, it usually can give rise to a feasible 1-bit counter.

**Figure 15:**
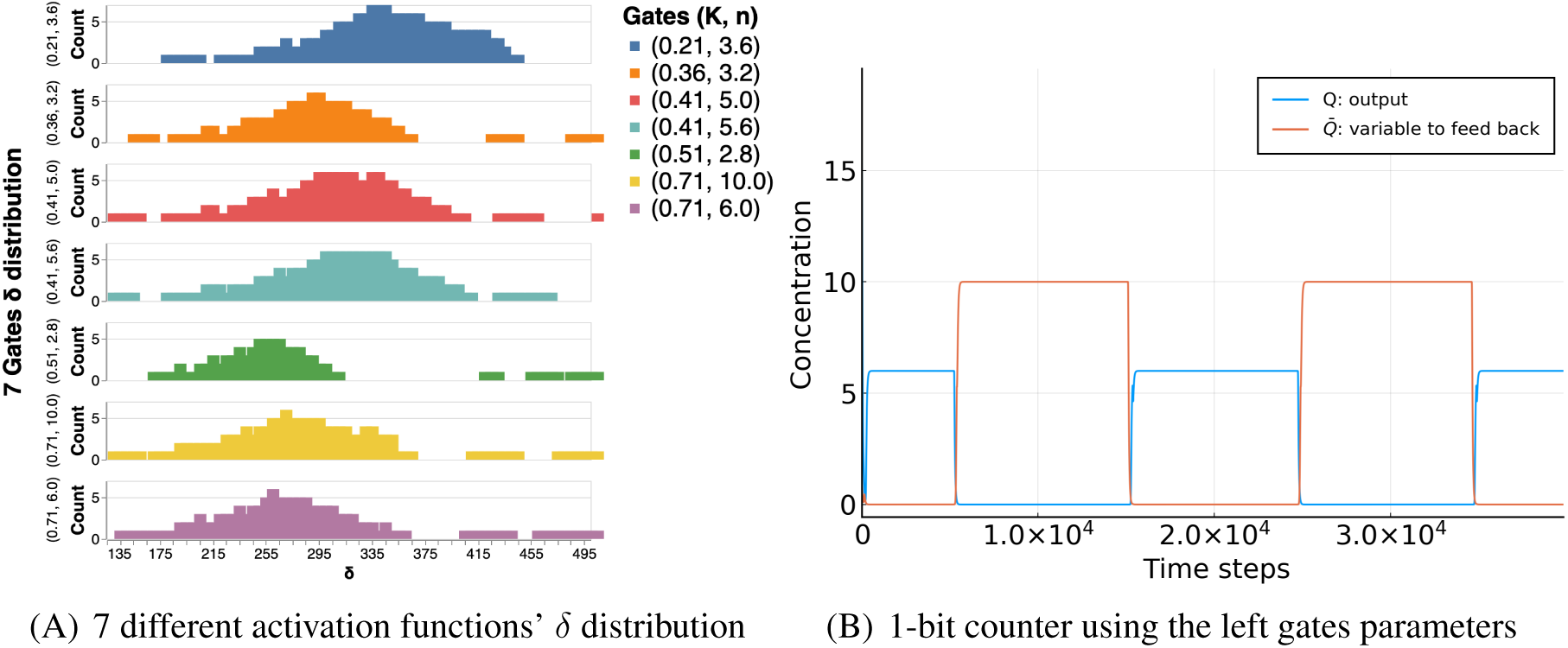
An example of a 1-bit counter with unique activation functions for each gate. A) Given an external pulse width *δ*, seven gates are selected conditioned on having the target pulse width within their distribution. Here we show the *δ* distributions associated with each gate that contains different (*n, K*). B) With the selected seven gates’ parameters from panel a), we show an example trajectory of a 1-bit counter.

Regarding the uniqueness of the optimal solution for a general *N* -bit counter, we quantify the optimality of an *N* -bit counter by *δ_cv_* which is the percentage of the coverage of the input pulse width within the counter’s maximum allowed *δ* range. For example, Supplementary Tables 1 and 2 list *δ_cv_* for various circuits. Also, we have generated the corresponding boxplot in Supplementary Figure 6 for a better visualization. Computationally, the optimal solution for a 1-bit counter is uniquely associated with a particular (*n, K*) pair. However, for multi-bit counters, there can exist multiple circuits with the same *δ_cv_* . This provides experimentalists with more options to accommodate constraints that are not captured by our modeling framework.

In order to achieve a feasible counter which operates according to a specific parameter set, one could perform a conditional sampling to generate a synthetic counter from the database. Using such a strategy for generating feasible 1-bit counters shows much greater flexibility in choosing a different desired range of activation functions. This affords more options to fine-tune the gates’ parameters and avoid construction constraints. Moreover, a machine learning classifier could be used to produce feasible 1-bit counter designs with an expected probability associated with each gate.

## 3 Discussion

Synthetic biology is an emergent interdisciplinary field of research, whose ultimate aim is to design novel biological systems, or to redesign existing ones, with the purpose of achieving new functionalities in medical, energy, environmental, chemical threat detection, and other applications (Cheng & Lu 2012, Andrianantoandro, Basu, Karig & Weiss 2006, Cameron, Bashor & Collins 2014), as well as contributing to the understanding of natural processes. In engineered synthetic circuits, basic cellular components such as genes, mRNAs, proteins, and metabolites are “rewired” in order to construct bio-molecular circuits that realize desired computations and capabilities (Sprinzak & Elowitz 2005). The resulting devices sense data, such as the presence or absence of particular surface features on a cell being interrogated, or of a chemical in the environment. They perform logical operations on this data and, based on the results of these operations, decide upon appropriate responses – such as the secretion of a toxic chemical in order to kill a cell that has been identified as being part of a tumor or as being infected by a pathogen, or the expression of a fluorescent signal to indicate an undesired trait. Specific application examples include the regulation of cell fate decisions (Sedlmayer, Aubel & Fussenegger 2018, Vogel, Persson, Seamons & Deans 2019), building cellular memory (Inniss & Silver 2013), engineering CAR-T cells for adaptive immunotherapy (Cho, Collins & Wong 2018), designing epigenetic reading and writing systems (Park, Patel, Keung & Khalil 2019), and developing numerous other specialized circuits (Callura, Cantor & Collins 2012, Archer, Robinson & Süel 2012, Elowitz & Leibler 2000, Yang, Nielsen, Fernandez-Rodriguez, McClune, Laub, Lu & Voigt 2014, Gupta, Bram & Weiss 2013, Hwang, Tan, Koh, Ho, Poh & Chang 2014, Saeidi, Wong, Lo, Nguyen, Ling, Leong, Poh & Chang 2011, Kohanski & Collins 2008).

One of the basic information-processing circuits is the sequential counter, which keeps track of the number of events of a certain type that have been detected. In any digital computer, precision is limited, and counts are always made modulo an integer; we take the same view here. We see the *N* -bit counter as a key component of future genetic computing devices, and analogous ideas should be useful in building more complicated feedback components. Potential applications of such a circuit would include programmable cell death after detecting a certain number of events (Khalil & Collins 2010). The resetting capability of the counter enables it to function periodically, such as directing the expression of specific enzymes in an ordered and periodic manner. It can also produce an output to match a natural cycle, for example based on sensed hormones and other circadian signals. Additionally, parity checkers, or more generally counting modulo an integer, are used in error-correcting codes, and can be expected to play an important role in synthetic digital inter-cellular communication devices.

Our work can be viewed in the broader context of modeling and mathematical and control-theoretic analysis in synthetic biology seen as an engineering discipline (Del Vecchio, Qian, Murray & Sontag 2018<u>b</u>). These analysis tools help guide bottom-up approaches to synthesis based on combining individual genetic components (Bloom, Winkler & Smolke 2014), modules (Hasty, Dolnik, Rottschäfer & Collins 2002), and programmed feedback (Bloom et al. 2014, Bloom, Winkler & Smolke 2015, Stapleton, Endo, Fujita, Hayashi, Takinoue, Saito & Inoue 2012). Novel engineering approaches and designs have greatly enhanced the ability to harness gene regulatory circuits for various applications (Bacchus, Aubel & Fussenegger 2013, Endo, Hayashi, Inoue & Saito 2013, Weber & Fussenegger 2009, Tigges, Marquez-Lago, Stelling & Fussenegger 2009), implementing Boolean logic functions (Miyamoto, Razavi, DeRose & Inoue 2013, Purcell & Lu 2014, Rinaudo, Bleris, Maddamsetti, Subramanian, Weiss & Benenson 2007, Daniel, Rubens, Sarpeshkar & Lu 2013, Farzadfard & Lu 2014, Roquet & Lu 2014, Bonnet, Yin, Ortiz, Subsoontorn & Endy 2013), and targeting specific disease states (Aubel & Fussenegger 2010).

In this work, we proposed a generic framework for building a distributed synthetic multi-bit counter. We are ultimately motivated by the long-term goal of designing synthetic genetic circuits that are “universal” computing devices; such circuits should be capable of finite-automaton computation, in analogy to central processing units in digital computers or, more abstractly, the “head” of a Turing machine. We started by designing the building block module, the single-bit counter (a parity checker). We then scaled-up the single-bit counter design to a multi-bit counter, distributing the computation using connector modules communicating between the individual parity checkers via diffusible small molecules. We have showcased a feasible single-bit counter design by implementing a Cello-optimized transcriptional circuit in the *E. coli* system *in silico*. Next, we have systematically explored the parameter space of genetic circuits with a shared shifted Hill activation function across all gates. The results can then be used to populate a database for designing more general counters. Our generic computational framework is universal, and the connector modules are flexible enough to be used with different cell-cell communication signaling systems (Fredriksson, Lagerström, Lundin & Schiöth 2003). For example, the communication signals can be realized not only by quorum-sensing molecules but also, potentially, by small peptides, and the EGFR system (Wieduwilt & Moasser 2008). Our multi-bit counter framework has quantifiable robustness against variation in the width of the input pulses, and it can naturally handle asynchronous counting situations. Our computational design framework can shed light on how experimentalists can build more robust distributed multi-bit counters by tuning the characteristic curves (described by Hill functions) of the gates to better match the range of expected pulse widths . This computational framework can significantly reduce the number of trial-and-error experiments by using the design selection graphs provided in Figure 14 as a showcase example.

Our work, while theoretical, lays out a clear path to the development of scalable counter. The experimental realization is feasible, as shown by the obtained designs and specific DNA sequences. On the other hand, the size of the required plasmids makes the actual experiments to be just beyond the edge of current technology. New tools (at Voigt’s lab and others) are being developed which will enable an implementation in the future.

Over the last decade, synthetic biology has played an increasing role in the understanding and controlling of complex cellular systems. However, numerous challenges remain. In order to connect different gates, one needs to deal with the problem of matching different dynamic ranges of multiple circuit outputs (Brophy & Voigt 2014), and synthetic circuits place a burden on host cells. Traditional genetic design was very labor-intensive, but this process has been greatly streamlined with the advent of genetic circuit design automation pipelines, including Cello (Yaman, Bhatia, Adler, Densmore & Beal 2012, Bhatia, Smanski, Voigt & Densmore 2017, Nielsen, Der, Shin, Vaidyanathan, Paralanov, Strychalski, Ross, Densmore & Voigt 2016<u>a</u>). It is now possible to obtain a promoter wiring diagram of a synthetic plasmid by supplying a Boolean function specified by an input-output truth table to Cello. Cello then attempts to minimize host burden while at the same time automatizing the search for optimal designs by employing simulated annealing and other optimization algorithms so as to minimize a “circuit score” with respect with a biological constraint library.

However, combinational logic is not sufficient. The intricate feedback wiring networks in living systems are evidence that cellular systems can dynamically perform complex jobs via memory-like units containing sequential logic circuits, and show how desired cell states emerge with cellular sensing. For example, living cells that differentiate into multi-cellular structures are usually controlled by sequential logic instead of purely combinational logic. One of the most compact examples for understanding cellular sequential logic is the “switch”-like memory system. By the turn of the millennium, Gardner et al. (2000) constructed a genetic toggle switch containing promoters that drive mutually inhibitory transcriptional repressors’ expression. This work has laid the foundation for building memory-based circuits, and it serves as a crucial framework for building the SR-latch used to realize the single-bit counter in our generic design. Later on, Friedland et al. (2009) studied two versions of genetic counter in *Escherichia coli* that can count up to three induction events only. In comparison, our 2-bit counter design has the advantage of counting an arbitrary number of pulses modulo 4.

Several synthetic circuits with memory have been proposed in the literature (Inniss & Silver 2013). For example, the work by Hillenbrand, Fritz & Gerland (2013) has studied a JK-latch. Instead of using the universal logic gate “NOR” everywhere, the authors proposed the JK-latch with a design combining “AND” gates and “NOR” gates. Such a design is not readily compatible with CRISPRi-based automation toolboxes (such as Cello), making it harder to implement and scale-up. In 2014, a genetic sequential logic circuit was explored with a clock pulse generator by Chuang & Lin (2014), thus relying upon periodicity of inputs. In contrast, our design naturally handles asynchronous counting scenarios. A counter design for a pulse-detecting circuit that only responds to the falling edge of the input pulse was proposed by Noman et al. (2016), detecting the completion of an event; in contrast, our design detects pulses with durations within a specific interval.

With the growing importance of engineering synthetic genetic circuits, we are currently facing scalability bottlenecks caused mainly by the inability of host cells to endure “large” foreign circuits. Due to the limitations of traditional recombinase-based state machines (Chiu & Jiang 2017, Yehl & Lu 2017), the CRISPRi system is being explored more actively to encode cellular-based memory (Yehl & Lu 2017, Andrews et al. 2018<u>b</u>). Our counter framework can be readily used with CRISPRi for implementing more complex circuits in host cells and can potentially realize additional functions per cell (Jusiak, Cleto, Perez-Piñera & Lu 2016). Al-though some designs for multi-cellular recombinase logic have been proposed lately (Guiziou, Ulliana, Moreau, Leclere & Bonnet 2018), a generic design framework with integrated feedback loops that performs asynchronous distributed counting has not been proposed so far.

CRISPRi holds the potential of being the right candidate for realizing more complex functions and offering more gates in a single cell. However, it critically relies on high expression levels of Cas9 which is toxic to the host cells (Zhang & Voigt 2018). Hence, adopting new principles and new synthetic architectures like distributed computation have also been widely discussed (Xiang, Dalchau & Wang 2018, Karkaria, Treloar, Barnes & Fedorec 2020, Al-Radhawi et al. 2020). Similarly, our approach for designing a multi-bit counter system tries to solve scalability issues via distributed computation using cell-cell communication. Until now, a generic scalable paradigm for implementing distributed computation with regulatory feedback for multi-cellular counting tasks had not been proposed. As a simple example for realizing distributed computation with memory in the sequential logic realm, our synthetic genetic counter can be widely adopted in various biological counting scenarios. The computational framework can be easily translated to be used in other biological contexts. The advantage of such a counting framework is not only restricted to being implemented as an event detection machine. It can be viewed as a thresholding machine for selecting sequential binding events in biology, such as in kinetic-proof reading systems (Hopfield 1974). Viewing the counter as a probing system can help us understand cellular mechanisms more deeply. It has been suggested that sequential logic can overcome the information bottlenecks inherent in complex networks (Letsou & Cai 2016). The combinatorial binding of transcription factors at promoters likely contributes to cell-type-specific gene expression in a static view. However, sequential logic may reveal a dynamic picture of the underlying reaction landscape.

As for future directions, we are intrigued by the idea of applying similar methods in mammalian cells, and specifically for immunotherapy. For example, one can think of engineering CAR-T cell systems for distributed immune system computation (using cytokines as signals such as in the work by Cho et al. (2018)), with the integrated ability to sense an external cancerous environment with tunable thresholding. Another example stems from developmental biology, where the distributed multi-bit counter can be considered as a guiding system for triggering various biochemical events at different developmental stages. Each bit level can be implemented to produce a specific set of transcription factors or essential proteins to trigger the desired cellular signaling events.

Furthermore, in another direction, epigenetic modifications play crucial roles in cell fate networks and the different developmental stages of a cell. A long-term goal in epigenetics is to realize systems that sense and count certain cellular events (Prochazka, Benenson & Zandstra 2017, Sheth & Wang 2018, Bashor & Collins 2018). A synthetic epigenetic system can be implemented with the ability to over-express TET to affect DNA methylation and other transcriptional factors to further affect RNA modifications, DNA methylation, and histone modifications. Promisingly, a very recent discovery (Hino, Rossetti, Maŕın-Llauradó, Aoki, Trepat, Matsuda & Hirashima 2020) of an ERK-mediated mechano-chemical feedback system that can generate complicated multi-cellular patterns has shed light on how cellular waves can contribute to the long-range correlation between different distant modules. Such long-range communication mechanisms will complement the diffusive nature of cell-cell communication signaling. They can be built on top of our counter system to achieve additional complex cellular functions and be used as an additional approach for realizing parallel cellular computation.

### Limitations of Study

From an analytical point of view, our analysis is based on the assumption that cells grow in a homogeneous medium; therefore time-delays due to diffusion and other spatial effects are assumed to be negligible. Growth of colonies on plates will have to be modeled with partial differential equations and the behavior of the circuit may depend on the geometry of the setting.

From an implementation viewpoint, scaling up the counter will require a larger number of orthogonal diffusible molecules than available at present. Current technology provides six such diffusible molecules (four types of quorum sensing signals based on AHL plus two additional ones as proposed by Du, Zhao, Zhang, Wang, Huang, Tian, Luo, Luo, Wang, Xiang et al. (2020), thus limiting the realizable counters to four bits.

## Author Contributions

Conceptualization: T.C, M.A, C.V, and E.S; methodology and investigation: T.C, M.A, and E.S; Formal analysis: T.C, M.A; Visualization: T.C; Data curation: T.C; writing – original draft, T.C; writing – review & editing: T.C, M.A, C.V, and E.S. Project administration: E.S; Funding acquisition: E.S.

## Acknowledgement

We thank Shuyi Zhang for initial discussions.

## Declarations of interests

The authors declare no competing or financial interests.

## 4 STAR *\** METHODS

### KEY RESOURCES TABLE

#### Lead contact

Further information should be directed to Eduardo Sontag

#### Materials availability

N/A

#### Data and code availability

The programming code that was used to analyze the raw data that supports the findings of this study is available in Github https://github.com/danielchen26/circuit-design.

### METHODS DETAILS

This work has proposed a new distributed circuit construction framework for a counter. We first show how we design the building block unit, which is a single-bit counter. Next, we describe our multi-bit counter system model that utilizes the same single-bit counter construction repetitively for scaling up the system. This section will first show the single-bit counter promoter design obtained by using available automated genetic design software (Nielsen et al. 2016<u>b</u>). We will explain how we come up with the gate assignment and perform the consequent mathematical modeling. All our simulations are performed using Julia (version 1.5.1) (Rackauckas & Nie 2017).

### 4.1 Biological gate design & assignment

#### Biological gate design

To show a working example for the single-bit counter in *E. coli*, we employ the genetic automation pipeline Cello 2.0 to map the logic gates to biological gates. The feedback in our system and the fact that inputs will be pulses introduce a major difficulty to an off-the-shelf application of Cello 2.0. This is since the current version of Cello handles combinational circuits only. Nevertheless, a recent extension of Cello was employed to design sequential logic circuits (Andrews et al. 2018<u>b</u>), and we use the latter version here.

In order to assign gates to our parity checkers, we consider a slight variation in which we temporarily ignore the feedback part. Thus, we first specify a state transition table (Table 1) for the open-loop circuit without feedback, and consider all possible inputs to the XNOR gate. The Cello 2.0 software takes this binary transition state table, together with the circuit topology specified by a Verilog file, and generates a biological gate assignment. Table 1 presents all possible transition states for the 2 inputs (*X, Y* ) and 2 outputs (**Q**, 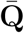) of the open-loop version. Cello 2.0 outputs a gate assignment that is optimized with respect to the cross-talk between different biological promoters, and aims at producing the best dynamical range of the overall circuit. However, Cello 2.0’s optimization algorithm does not maximize the range of *δ* (pulse duration), which is the relevant objective function in our design. Hence, we have performed an additional optimization step by replacing one gate at a time with another gate from the *E.coli* gate library. We tested all such one-gate-replacement candidate circuits and compared them to the Cello-produced circuit. The optimized circuit is depicted in supplementary Figure 10 (B) which shows the biological gate diagram of the plasmid based on the following repressors: QacR, LmrA, SrpR, BM3R1, PhIF, PsrA, BetI. This diagram follows Cello conventions and has produced the best *δ* range among all candidates. The list of all gates is given in Table 2. In the SI, we show other feasible circuits.

**Table 2:** Gate number - biological gate mapping

| gate number | biological gate |
| --- | --- |
| 1 | QacR |
| 2 | LmrA |
| 3 | SrpR |
| 4 | BM3R1 |
| 5 | PhIF |
| 6 | PsrA |
| 7 | BetI |

The diagram in supplementary Figure 10 (B) is based on the topology that was specified by our open-loop circuit Verilog file. However, our desired design contains a feedback path from the BetI gate to the QacR gate. How does one biologically “connect” this output signal back to the circuit’s input to obtain the closed-loop version of the circuit? The “gene-promoter” paradigm is perfectly suited to achieve such a goal: the answer is to copy the promoter in front of the PsrA gene (which is the target of the BetI gene as part of the SR-latch) to the promoter region of the QacR gene (which is a tandem promoter also containing the target of the input signal, pBAD). Thus, the QacR gate will receive a signal from the BetI gate without changing the characteristic response curve for the gate. Changes in the promoter region of a gate do not change the response curve of that particular gate because the parameters that define the response curve only relate to what is inside the “gate box”, namely the gene coding region and the promoter region of a target gene that the protein expressed by the previous gene can bind to repress.

#### Mathematical model for the single-bit counter

As the repressors have been assigned to the logical gates, we introduce a differential equation model and use it to analyze the effects of closing the feedback loop and applying the input pulses.

Using the gate assignments from Cello 2.0, we are now able to write down our differential equation model for the single-bit counter. In this case, pBAD is considered as the external input, and the BetI gate feeds back to the QacR gate. The circuit output will be the yellow fluorescent protein (YFP) produced by the gate PsrA.

Our model is given by the following system of seven differential equations:

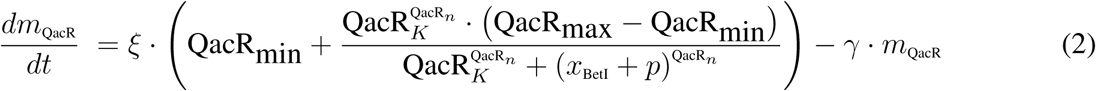

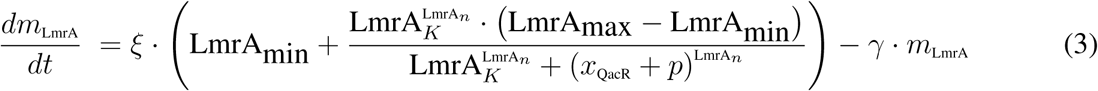

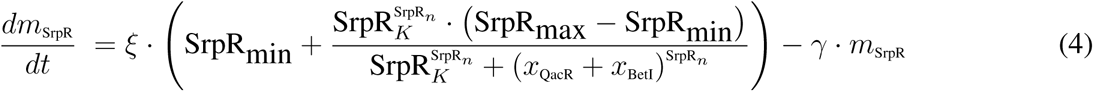

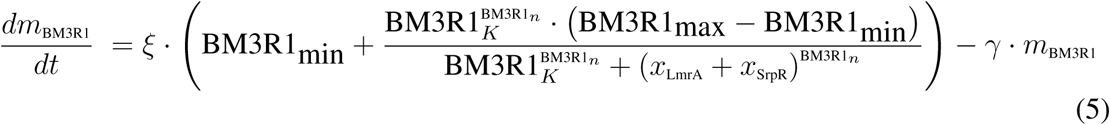

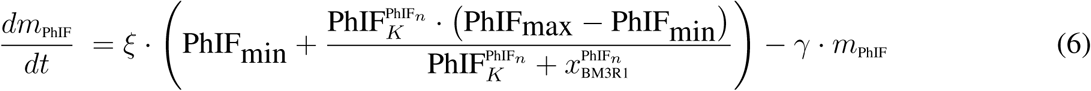

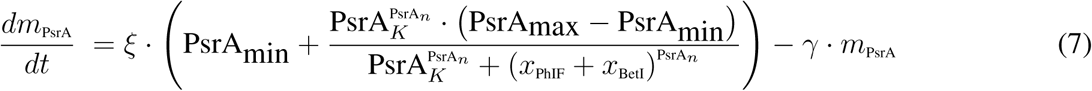

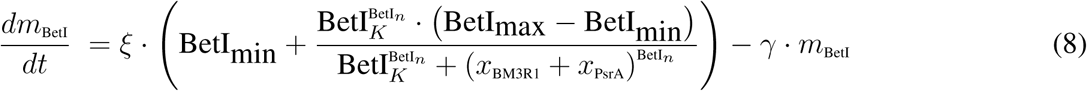

This model, and its parameters, are derived from the Cello 2.0 specifications as well as the paper by Andrews et al. (2018<u>a</u>). The differential equation model has seven state variables *m_gate name_*, which represent the mRNA concentrations associated with the respective gates. (Protein concentrations are assumed to be proportional to these.) Within the parentheses, we model each gate with a Hill equation in terms of each gate’s relative promoter strength, and then convert units to mRNA concentrations by the scaling factor *ξ* . For each differential equation, we also include a degradation term for mRNA. For more details on this procedure, and the parameters used, see the work of Nielsen et al. (2016<u>b</u>).

The assumption made in such models is that mRNA production rate is proportional to the relative promoter units (RPUs) for each promoter’s activity. The *m_gate name_* is the concentration of mRNA produced by the output promoter. The constant *ξ* is a scaling factor that is used to convert relative promoter strength to mRNA concentrations for gate variables, and *γ* is the degradation rate of the mRNA of each gene. The promoter activities (7 gates) are reported in relative promoter units (RPUs). The input for each gate *x*_gate name_ = *ξγ^−^*^1^*m*_gate name_, noting that *ξγ^−^*^1^ = 1 for our choice of parameters. Both *ξ* and *γ* are taken as constants here. They were arbitrarily set to *γ* = 0.025 min*^−^*^1^ and *ξ* = 0.025 [mRNA]*/*min-RPU in order to produce time trajectories consistent with bio-molecular timescales (Andrews et al. 2018<u>a</u>).

### 4.2 Multi-bit counter

In the single-bit counter analysis, we have used the gates’ assignment from the *E. coli* system. Each biological gate is unique and has different activation Hill functions. Next, we show how we model the multi-bit counter in which all gates share a single activation function. This means that the parameter sets for each gate are the same across seven differential equations as shown above. The shared version differential equations are indicated in equation (9).

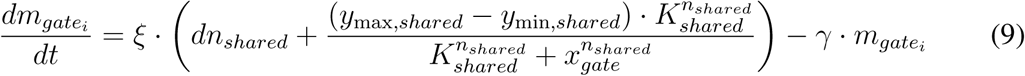

Since the multi-bit counter system is modular, for a counting system that counts to 2*^N^* , we will need *N* − 1 connectors, with a connector type 1 placed between the first two bits, and the rest of the connectors are all type 2. The dynamics of these two types of connectors are shown below.

#### Mathematical model for the Connector type 1

The connector type 1 (AND gate) module in Figure 8 can be modeled as follows. There are three gates in this module. Each gate’s dynamics is modeled by a shifted Hill activation function and a degradation function in general. The gate A and gate B in Figure 8 are two NOT gates with a single input, which is modeled by Eq. (10) and (11). The gate C is a NOR gate that takes the output from gate A and gate B as inputs, and the dynamical equation for gate C is given by equation (12). The overall system can be written as follows:

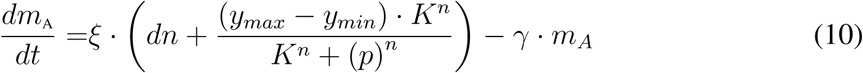

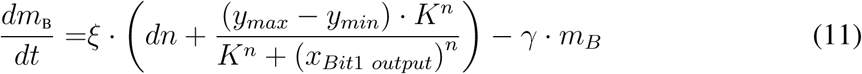

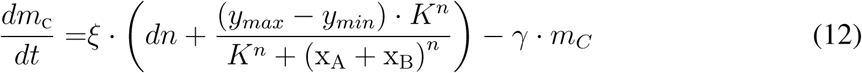

In this set of equations, the scaling factor (*ξ*) and the degradation rate (*γ*) are constants, which are the ones in the single-bit counter. **The shifted Hill function is specified by the parameters:** *y_min_*, *y_max_*, *K*, *n*. The meaning of the variables has been illustrated in Figure 3.

#### Mathematical model for the Connector type 2

With the diagram of the connector type 2 shown in Figure 8 (B). We use the following dynamical equations to model the dynamics of the 6 gates in the connector type 2 circuit module. In general, take the *n*-bit counter as an example, the connector between *n*th and the (*n−* 1)th 1-bit counters follow the following ODE:

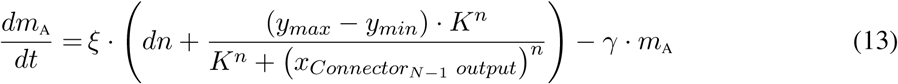

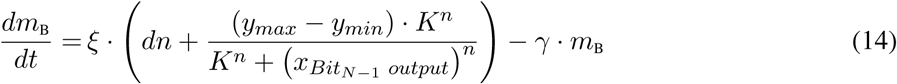

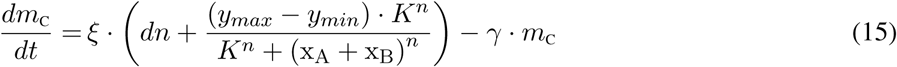

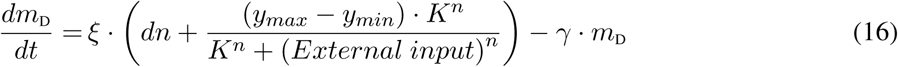

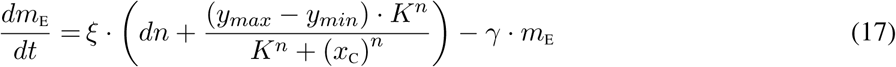

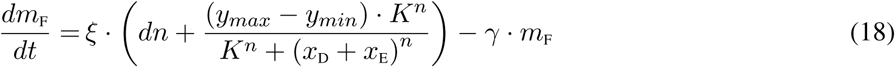

The parameters *ξ* and *γ* are constants and are the same as the ones described in connector type 1 dynamical equations. The shifted Hill coefficients are *y_min_*, *y_max_*, *K*, *n*.

So far, we have written all ODEs that describe the underlying dynamical systems for each of the modules. Therefore, one can build a full mathematical model for an arbitrary multi-bit counter just by merely assembling together the corresponding modules. Additionally, one should note that because we have assumed that repressor-promoter binding/unbinding dynamics occur at a fast time scale, we can assume that the promoter concentration (strain density) is at a quasi-equilibrium state. Next, we construct the counter’s parameter searching algorithm.

### 4.3 Parameter optimization

We optimize the parameters by a computational search. The goal is to find the valid range of the input pulse width and find the minimum duration for the relaxation time (defined as the time difference between the previous signal off time and the next input signal). Our simulation results show that within the valid range of the input duration and relaxation times, the designed circuits behave as desired.

The key to successfully constructing a feasible counter circuit is to find valid ranges of the parameters that the counting dynamics allows. Specifically, when an input signal is given to the counter, the SR-latch should exhibit a switching behavior. The appropriate duration of an input pulse is the one that makes the state of the SR-latch switch to the other steady state as shown in, Table 1.

All gates in the multi-bit counter system share a single input-output characteristic for the gate activation function. In reality, the activation functions for different transcriptional logic NOR gates can be different. Experimentalists have built a library of transcriptional gates with various dynamical ranges for different genetic gates. However, in our multi-bit counter design, we constrain all genetic gates to share the same activation function to reduce computational complexity and avoid combinatorial explosion. Feasible parameter ranges for the activation function will be explored for a multi-bit counter system.

The parameters essential for constructing a feasible multi-bit counter system are the shifted Hill coefficients *y_min_*, *y_max_*, *K*, *n*, the signal amplitude *A*, and the pulse width *δ*. The shifted Hill coefficients define the counter system’s intrinsic dynamics. The external signal amplitude and the pulse width can be controlled exogenously. From the engineering and control perspective, given a system dynamics with a fixed set of parameters for the shifted Hill coefficients, researchers will want to find the feasible range for the counter’s pulse width and amplitude. On the other hand, if researchers have an interest in a particular range of pulse widths, our framework will provide estimates of the optimal parameters.

In what follows, we include a detailed description of what the essential parameters are and why a feasible counter system depends on them:

- **External pulse width** *δ*. It specifies the duration of the external pulse. The feasible ranges of *δ*’s mean and standard deviation values are important indicators that represent the counter’s desired operating regime and the robustness against perturbations respectively.
- **External signal amplitude** *A*. It specifies the external signal strength. Knowing the feasible range for these parameters can be beneficial for experiments when dealing with both the quantity and cost of using chemicals or biological inducers.
- **The maximum value** *y_max_* **of Hill function.** To control it from an engineering perspective, one can manipulate the gates’ dynamical range to be reactive to certain input pulse widths. It also relates to the value of the circuit steady-state outputs. Controlling “up” can be useful when considering the counter system as a thresholding machine.
- **Equilibrium constant** *K*. It controls the cut-off point of the activation function.
- **cooperativity index** *n*. It controls the steepness of the activation function. In general, the synthetic gates with smaller “*n*” values turn out to be easier to be synthesized (Moon, Lou, Tamsir, Stanton & Voigt 2012).

In addition, if the characteristics of the gates are given, we need to determine the parameter ranges that are most robust to external perturbations. Specifically, for each group of Hill coefficient pairs (*n, K*), we need to determine the corresponding ranges of *δ*, *A* and *y_max_* around their means.

To determine if a counter is “good” or “bad”, we use a cost function to select the parameters. The general optimization procedure is stated as below:

- **Alignment of the carry bits:** For the 2-bit counter which is a special case of a general *N* -bit (*N >* 3) , the carry bit dynamics alone is an indicator of whether the connector type 1 can pass the information to the next level. Taking the 3-bit counter as an example, we consider that counting will start at the time when the two carry bits are first aligned, which coincides with the first time that the last bit switches, and thus sets the initial time reference point of the 3-bit counter dynamics.
- **Building a local switch cost:** In response to an external input signal, a local switch cost is needed to determine whether to switch the steady states at each bit level. For example, for the 1st bit level in the multi-bit counter system, one is expected to have the output switch from a low state to a high state (or reversed direction) upon detecting an input pulse. At the 2nd bit level, the output is expected to switch once every time the 1st-bit output switches twice. For the 3rd bit level, the output should switch once every time the 1st-bit output switches four times.
- **Periodic activation criterion:** For a multi-bit counter, especially when *N* is large, we require the first-bit to switch upon every input pulse stably. For example, a full cycle for a 3-bit counter requires at least eight stable switches for the 1st bit, four stable switches for the 2nd bit, and one stable switch for the 3rd bit. We have illustrated the switching pattern of the 3-bit counter in Figure 10 (B).
- **Global cost function:** In order to have a functional counter with a pulse that has the right duration, we need to combine all the constraints mentioned above for constructing a global cost function. In addition to the local cost criterion, we need to compute two additional quantities that can impose the conditions on combining all local switching costs at different bit levels altogether. These two quantities are:

First,

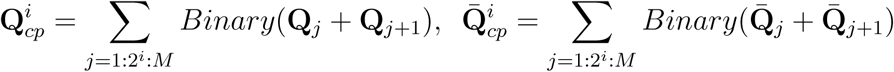

where *i* indicates the *i*th bit level for the multi-bit counter, *j* represents the index of the *j*th pulse and *M* represents the total number of input pulses seen by the counter. The notation *j* = 1 : 2*^i^* : *M* means to sum *j* from 1 to *M* every 2*^i^*. The “Binary” operator in the formula will compare the **Q** or 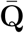 with *up/*2. if **Q** or 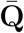 *> up/*2, the operator will return 1, otherwise it will return 0. Take the 3-bit counter as an example, for a full counting cycle in which the 1st bit takes 8 pulses, the 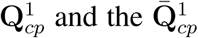 should be 4 for the 1st bit level, 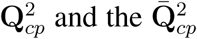 should be 2 for the 2nd bit level and 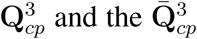 should be 1 for the 3rd bit level.

Second,

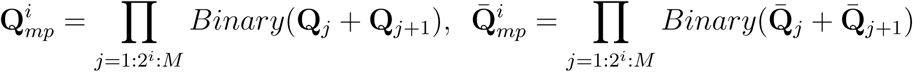

where *i* indicates the *i*th bit level for the multi-bit counters, *j* represents the index of the *j*th pulse and *M* represents the total number of input pulses that are inputs to the counter. The “Binary” operator has been defined above. If a set of parameter gives a feasible counter, then 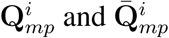 should all be equal to 0.

Therefore, when we impose the global condition for a feasible counter, we merge all constraints mentioned above with the “AND” operation for determining the feasible parameter set of a multi-bit counter. We define the overall cost function as follows:

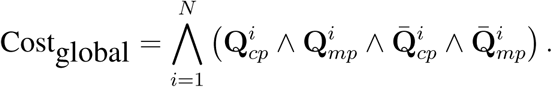

If all conditions are True, then Cost_global_ = 1, otherwise Cost_global_ = 0. With the cost function defined above, one can set the *N* -bit counter criterion with *M* pulses seen by the system. The above optimization will lead us to find the feasible parameter sets.

We summarize the procedure in the following algorithm. The pseudo-code algorithm shows how we optimize the system and find the feasible parameter ranges below:

**Algorithm 1.**
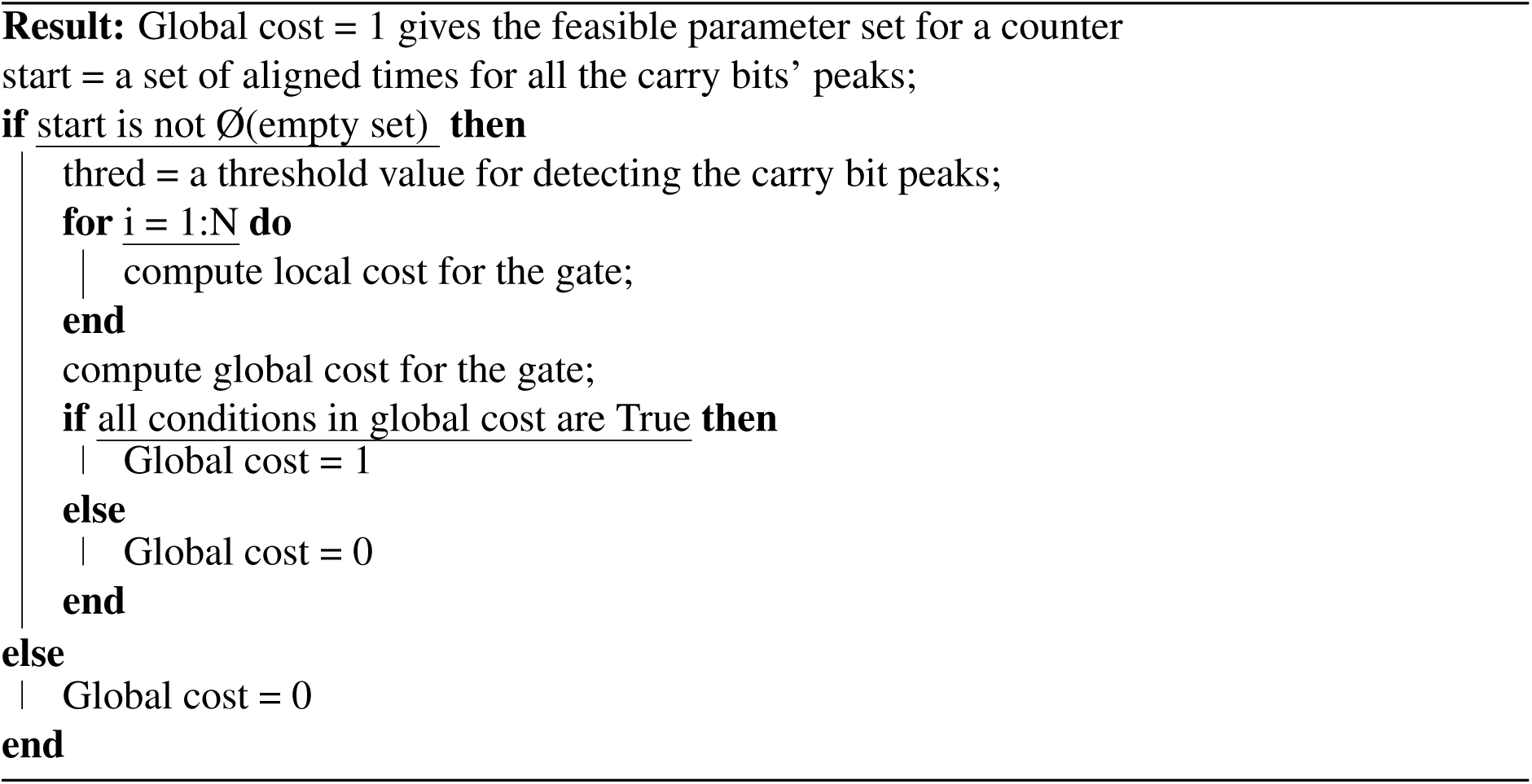
Optimization of choosing feasible parameters for a generic multi-bit counter.

With the general algorithm stated above, we can generate a database of feasible parameters for any multi-bit counter system. We showcase a snapshot of how a 3-bit counter database looks like in Figure 16. The parameter ranges are chosen according to the available experimental genetic gates database shown in Figure 11. Surprisingly, we found that *A* behaves mostly like a “free” variable that only needs to pass a low threshold value. In other words, *A* has a wide feasible range. This gives us the advantage of using a relatively small number of inducers for making the counter to work, and it also indicates that the counter can be less sensitive to the input signal concentration as long as it passes a small threshold.

**Figure 16:**
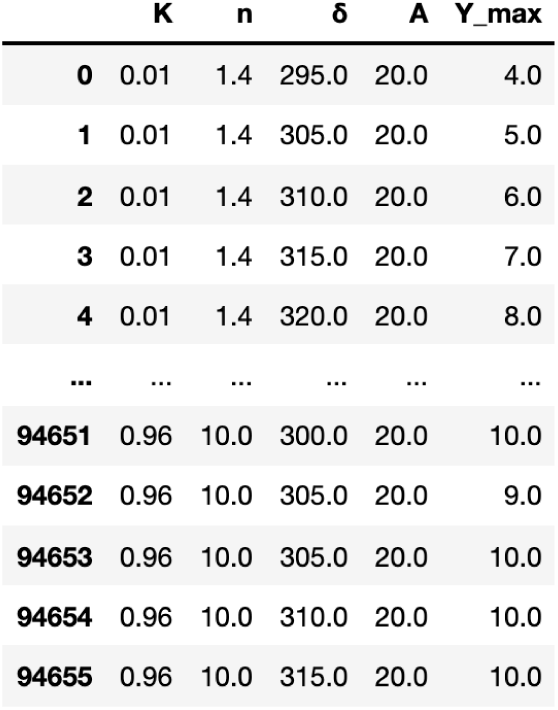
A snapshot of the database of the 3-bit counter. Each row in the data table corresponds to a feasible parameter set that gives rise to a working 3-bit counter. In this table, we only show what resulting database looks like, and the sampled parameter *K* ranges from 0.011 to 0.091, the parameter *n* ranges from 1.5 to 3.0, the parameter *δ* ranges from 275 to 350, the parameter *y_max_* ranges from 1 to 3, and we fixed the parameter *A* to 20. The full table can be found in https://github.com/danielchen26/circuit-design

## 5 Data availability

N/A

## Supplementary Information

### 1 Long-term behavior of the autonomous one-bit counter

In the absence of an external input, the 1-bit counter depicted in Fig. 2-A is a dynamical system capable of exhibiting multi-stationarity for certain parameter ranges. But in order to show that it is indeed a multi-stable system, we need to prove that every trajectory converges to a steady state. To that end, we can use known results [1] to show that the autonomous 1-bit counter is a monotone system that enjoys generic convergence to steady-states under mild conditions. This precludes the possibility of oscillatory or chaotic behaviors. An informal proof can be achieved by examining the influence graph of the network and verifying that every single loop has a positive sign as can be seen in Supplementary Figure 1. More details on the specifics of the method can be found in [1].

**Supplementary Figure 1:**
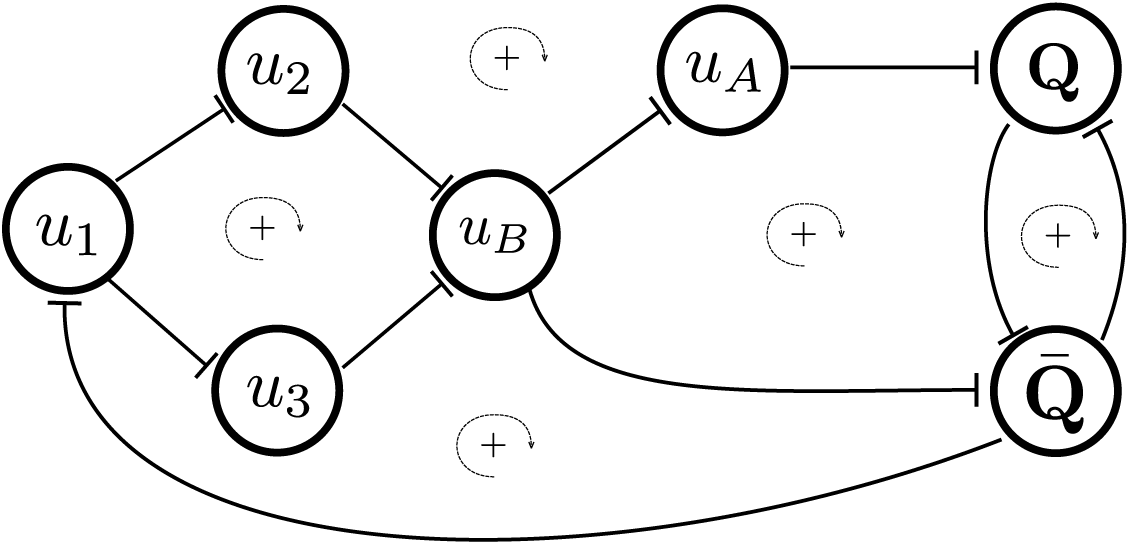
**Graphical test of monotonicity.** Related to Figure 2. Every (undirected) loop in the influence graph of the autonomous 1-bit counter has a positive sign. Every edge of the form “*f*” is assigned a negative sign. The sign of a loop is the product of the signs of the constituent edges.

### 2 Parameters that give rise to bistability and oscillation

In the main paper, we have discussed how bistability and oscillatory dynamics arise when varying the parameters *K*, *n*, and *y*_max_. For each point in the 3D parameter space, we evaluate if the corresponding parameter set produces multiple steady states for the 1-bit counter. This is accomplished by computing the trajectories corresponding to 100 random initial conditions to evaluate the number of distinct steady state solutions in the absence of an external input signal. Similarly, we find the parameter sets that allows the 1-bit counter to exhibit an oscillatory behavior when a constant input signal is applied. Combining these two numerical studies, we partition the 3D phase space into three distinct regions based on our simulation results. Supplementary Figure 2-(a) marks the regions “not bistable and not oscillatory”, “bistable and not oscillatory” and “bistable and oscillatory”. We select our parameters from the last region which has the potential of producing a functional 1-bit counter.

**Supplementary Figure 2:**
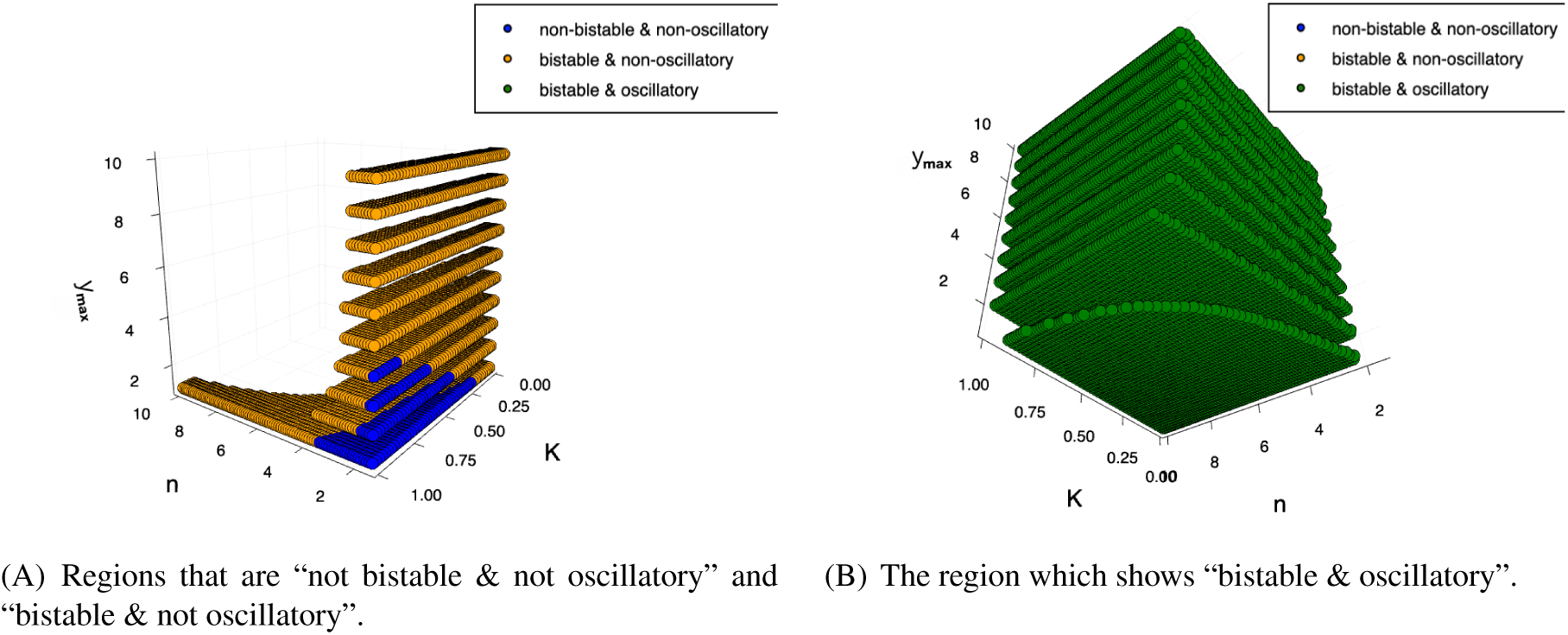
**3D bifurcation plots**. Related to Figure 5. (A) The layered 3D scatter plot shows which parameters give rise to “not bistable and not oscillatiory” or ”bistable and not oscillatory” systems. One can see a trend that as *y*_max_ goes up, the blue region becomes smaller and smaller. (B) This layered 3D scatter plot shows the parameter sets that correspond to “bistable and oscillatory”. We notice that as *y*_max_ becomes higher, the corresponding region in the space becomes larger.

### 3 Dynamics at the transition

In the main paper, Figures 9 and 10 show examples of the dynamics of the 2-bit and 3-bit counters. We indicate the external input pulses with green dots. However, due to the long relaxation times between consecutive pulses, the two green dots indicating the start and the end of a pulse can hardly be distinguished. Supplementary Figure 4 shows a zoom-in inset figure that clarifies the dynamics around the input pulse.

### 4 The *N* -bit counter (*N* = 8)

The scalability of a distributed system is an essential aspect when constructing larger circuits. We have shown systematically how to build a single bit counter, a two bit counter and a 3-bit counter in the main

**Supplementary Figure 3:**
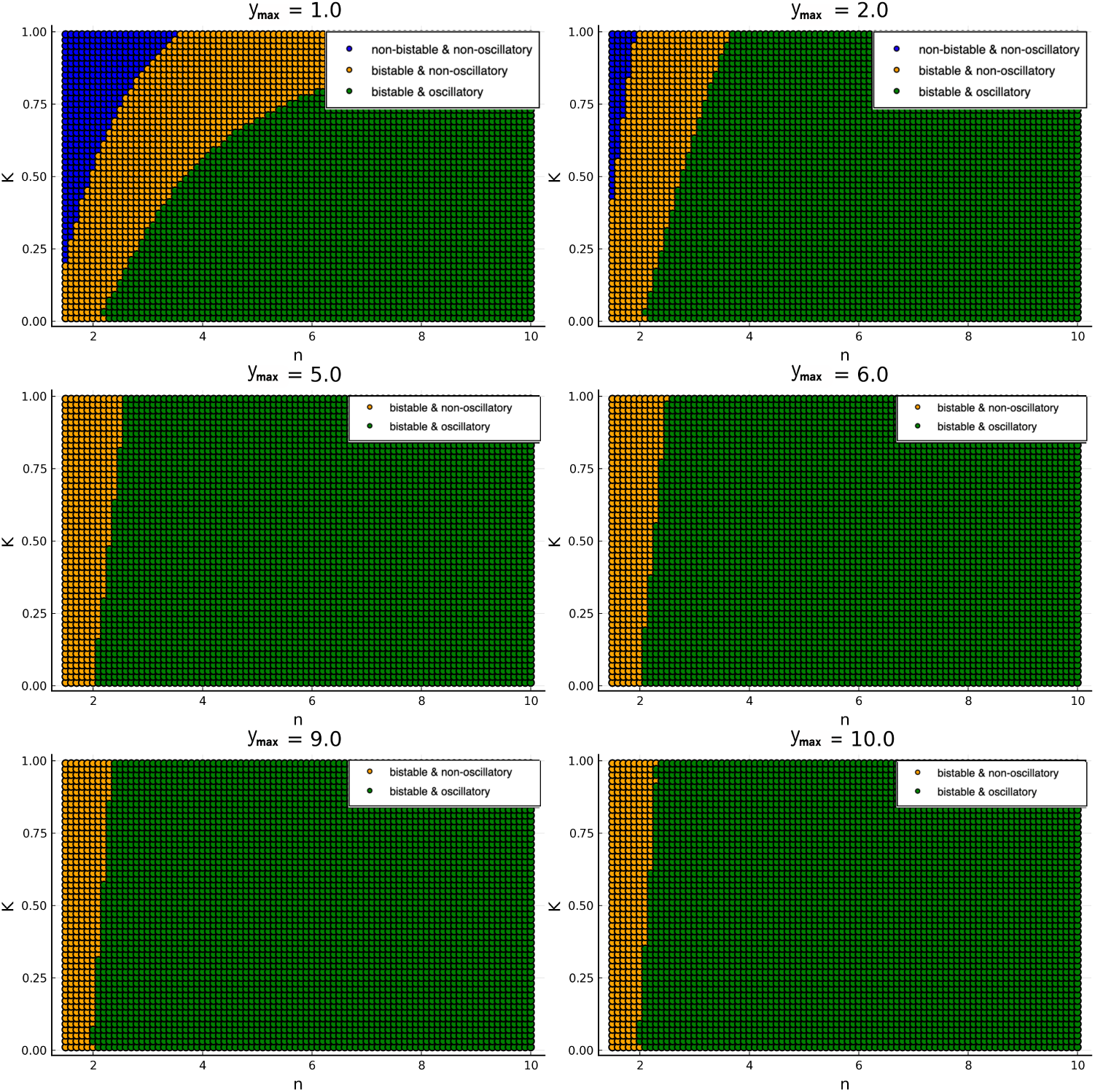
**A detailed layer-by-layer plot of the 3D bifurcation plot**. Related to Figure 5. paper. In order to demonestrate the scalability of the design, we design an 8-bit counter using the same principle. We show the simulation in Supplementary Figure 5 which demonstrates that all the bits switch as required.

### 5 Single-bit counter circuits implemented with seven unique gates in *E. coli*

In the main paper, we have shown an example of a single-bit counter design associated with an optimized range of the input signal duration *δ*. However, the Cello pipeline did not directly generate it. For our designed counter, Cello aims at generating the best candidate circuit with respect to several criteria. However, Cello does not take the range of *δ* into consideration. Here, we list seven feasible 1-bit counter circuits, each of which consists of seven unique repressors sampled from the *E.coli* experimental gate database.

Table 1 shows a comparison between the ranges of *δ* for these seven circuits. Each circuit is implemented with a unique set of seven gates sampled from the *E.coli* database. Circuit No. 1 in Table 1 was generated using the modified Cell 2.0 pipeline. We then changed one gate at a time in circuit No. 1, and replaced it with a gate sampled from the remaining gate pool in the *E.coli* database. We extensively tested all 42 possible combinations of gate replacements to search for a feasible counter. This has resulted in only 7 feasible designs. Note that *δ_range_* and *δ_cv_* of circuit No. 4 are significantly higher than those of circuit No. 1. This is not surprising since Cello generates a feasible set of gate assignments, but does not optimize *δ_range_*. Furthermore, to better visualize the distribution of *δ*, we show a box plot in Figure 6 for the seven circuits. Supplementary Figures 7-13 display the circuit plasmids, the gate assignments, and examples of the circuit dynamics with *δ* = 400 for each of the seven circuits.

**Supplementary Figure 4:**
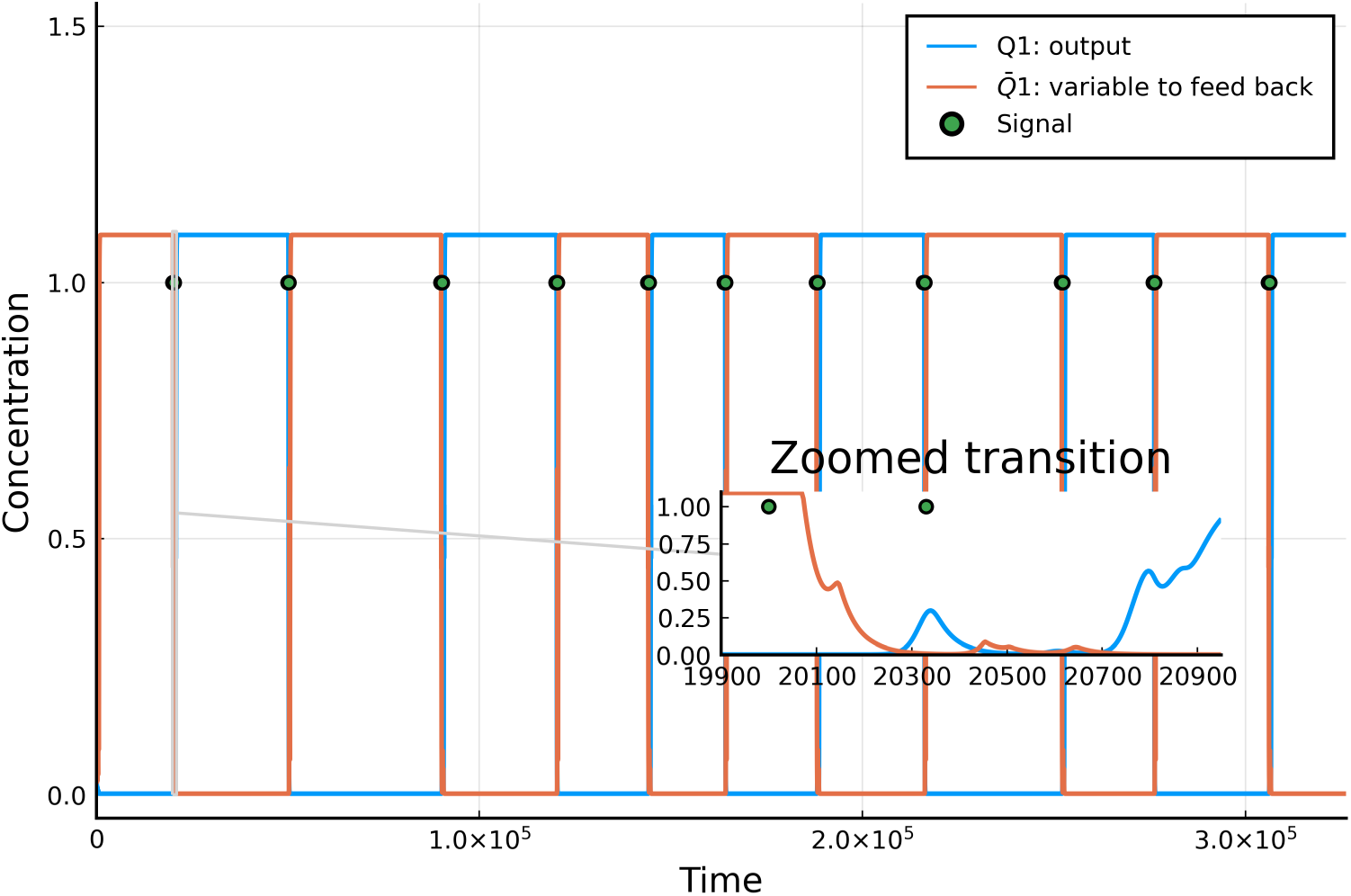
**Transition dynamics in the 1-bit counter**. Related to Figures 9 & 10. A inset plot for the dynamics in the interval *t* [19900, 20900] is shown in the right lower corner. When zoomed in, one can clearly see that there are actually two green dots indicating the start and the end of the corresponding pulse.

### 6 Contour plots

In the main paper, Figure 16 shows a comparison between *δ*-mean and *δ*-std for both the 1-bit and the 2-bit counters. Here we show an alternative representation as contour plots in Figure 14. These plots allow one to see how the *δ*-mean and the *δ*-std change when the number of bits in the counter are increased.

---

**Table 1:**
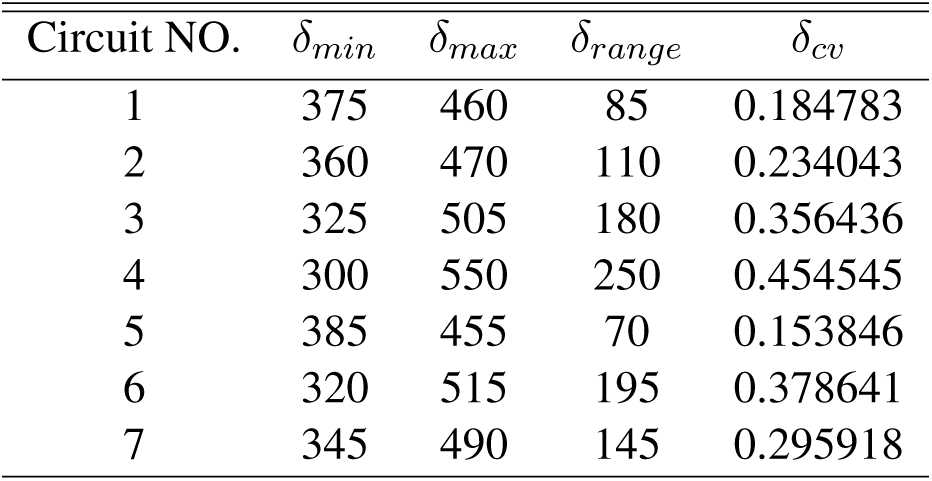
Comparison between the ranges of *δ* for the seven single-bit counter circuits. Each circuit is implemented with a unique set of seven gates from the *E.coli* database. *δ_min_* is the minimum value of *δ* to produce a feasible 1-bit counter, while *δ_max_* is the maximum value of *δ* that allows for building a feasible 1-bit counter. *δ_range_* is the difference between *δ_max_* and *δ_min_*. *δ_cv_* tells the percentage of the coverage for the input signal duration within the counter’s maximum allowed *δ* range.

| Circuit NO. | $\delta_{min}$ | $\delta_{max}$ | $\delta_{range}$ | $\delta_{cv}$ |
| --- | --- | --- | --- | --- |
| 1 | 375 | 460 | 85 | 0.184783 |
| 2 | 360 | 470 | 110 | 0.234043 |
| 3 | 325 | 505 | 180 | 0.356436 |
| 4 | 300 | 550 | 250 | 0.454545 |
| 5 | 385 | 455 | 70 | 0.153846 |
| 6 | 320 | 515 | 195 | 0.378641 |
| 7 | 345 | 490 | 145 | 0.295918 |

**Table 2:** The gate assignments for the seven circuits. To match to Cello 2.0 style, we have concatenated the associated RBS name (the first 2 letters of the gate name) with each gate name as a whole.

| Circuit NO. | Gate1 | Gate2 | Gate3 | Gate4 | Gate5 | Gate6 | Gate7 |
| --- | --- | --- | --- | --- | --- | --- | --- |
| 1 | H1_HlyIIR | N1_LmrA | S4_SrpR | B3_BM3R1 | P1_PhIF | R1_PsrA | E1_BetI |
| 2 | H1_HlyIIR | F1_AmeR | S4_SrpR | B3_BM3R1 | P1_PhIF | R1_PsrA | E1_BetI |
| 3 | H1_HlyIIR | N1_LmrA | S4_SrpR | B3_BM3R1 | Q1_QacR | R1_PsrA | E1_BetI |
| 4 | Q2_QacR | N1_LmrA | S4_SrpR | B3_BM3R1 | P1_PhIF | R1_PsrA | E1_BetI |
| 5 | H1_HlyIIR | N1_LmrA | S4_SrpR | B3_BM3R1 | P1_PhIF | R1_PsrA | Q2_QacR |
| 6 | A1_AmtR | N1_LmrA | S4_SrpR | B3_BM3R1 | P1_PhIF | R1_PsrA | E1_BetI |
| 7 | L1_LitR | N1_LmrA | S4_SrpR | B3_BM3R1 | P1_PhIF | R1_PsrA | E1_BetI |

**Supplementary Figure 5:**
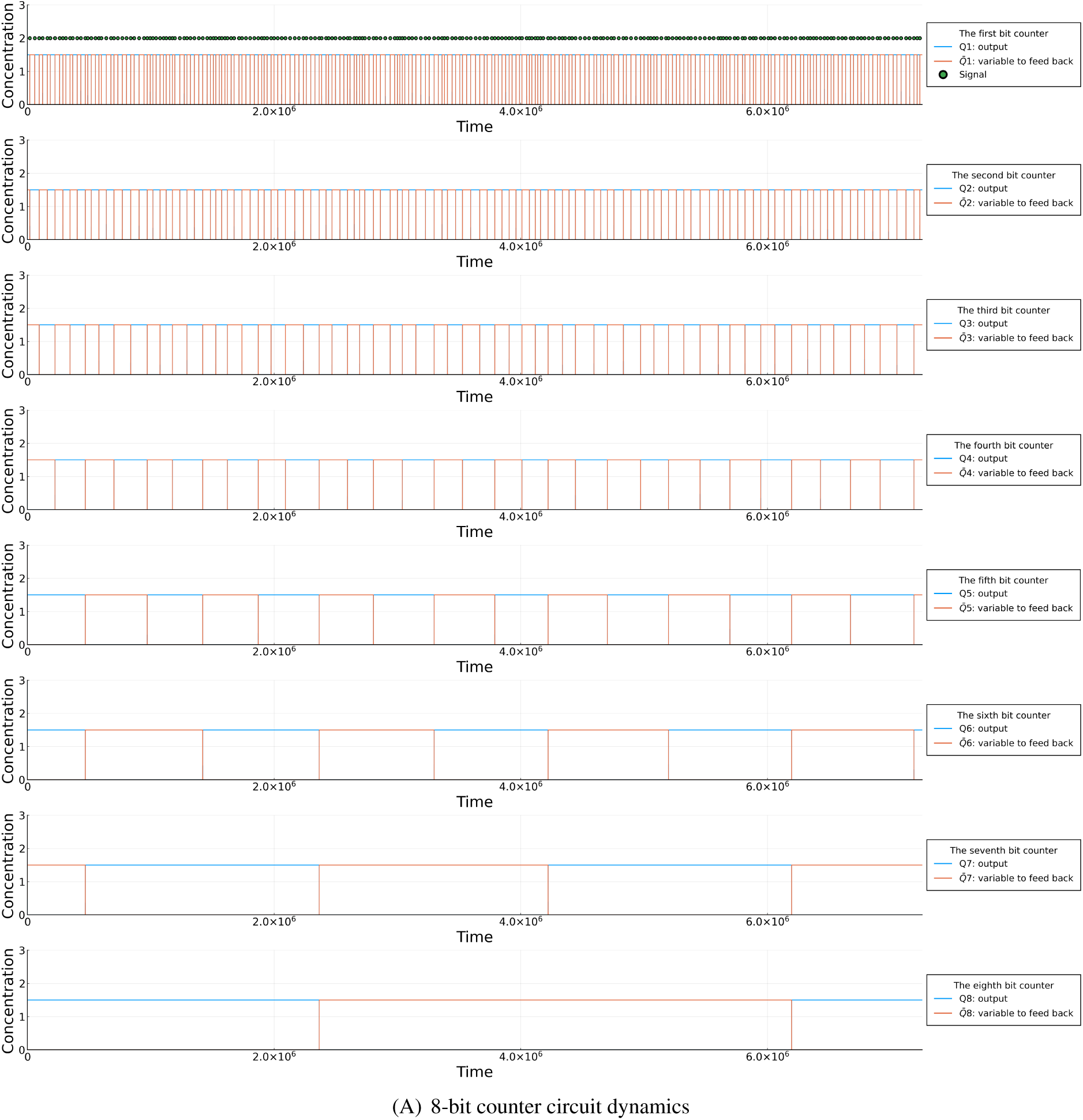
**Simulation of the dynamics of an 8-bit counter**. Related to Figure 6. The bits are ordered from the least significant bit to the most significant bit. The top panel is the first bit and the green dots indicate the input signals. The parameters are: the scaling factor *ξ* = 0.025, the degradation rate *γ* = 0.025, the equilibrium constant *K* = 0.081, the cooperativity coefficient *n* = 2.81, the duration of external signal *δ* = 270, the amplitude of the external signal *A* = 20, the shifted hill coefficient maximum value *y_max_* = 1.5, the shifted hill coefficient minimum value *y_min_* = 0.002, and the initial and between signals relaxation times Δ = 20000. Note that the pulses are applied at random non-uniform intervals, which represents an asynchronous counting situation. However, the spacing between the bits looks more uniform for the more significant bits. This can be explained as follows. Suppose that the pulses are separated by random independent and identically distributed inter-arrival times; by the central limit theorem, the net effect of*_√_N* pulses will be distributed with a Gaussian distribution that has a variance which is proportional to 1*/ N* .

**Supplementary Figure 6:**
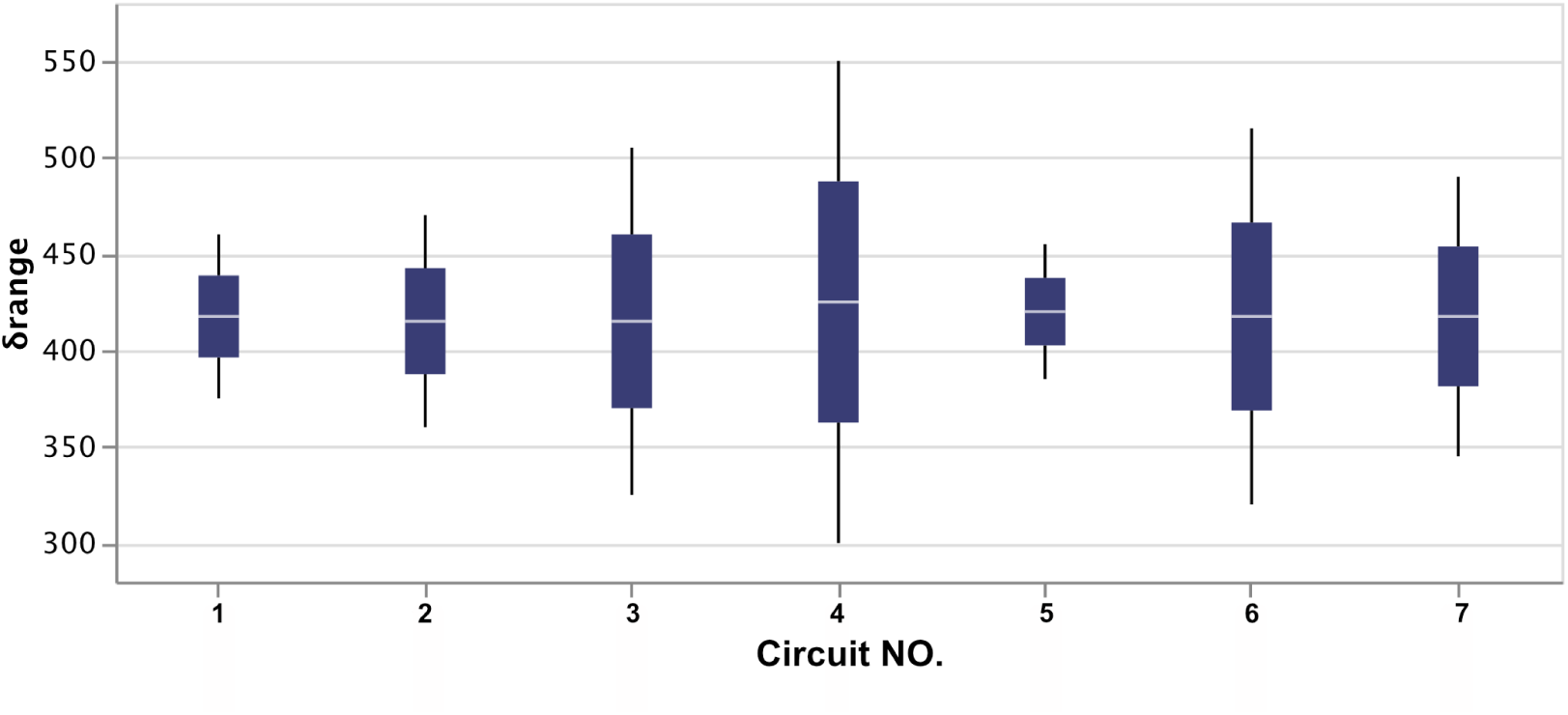
**The distribution and the feasible ranges of *δ* for all the seven 1-bit counter circuits**. Related to Figure 2 & 15. Note that circuit No. 4 has the largest range of *δ*.

**Supplementary Figure 7:**
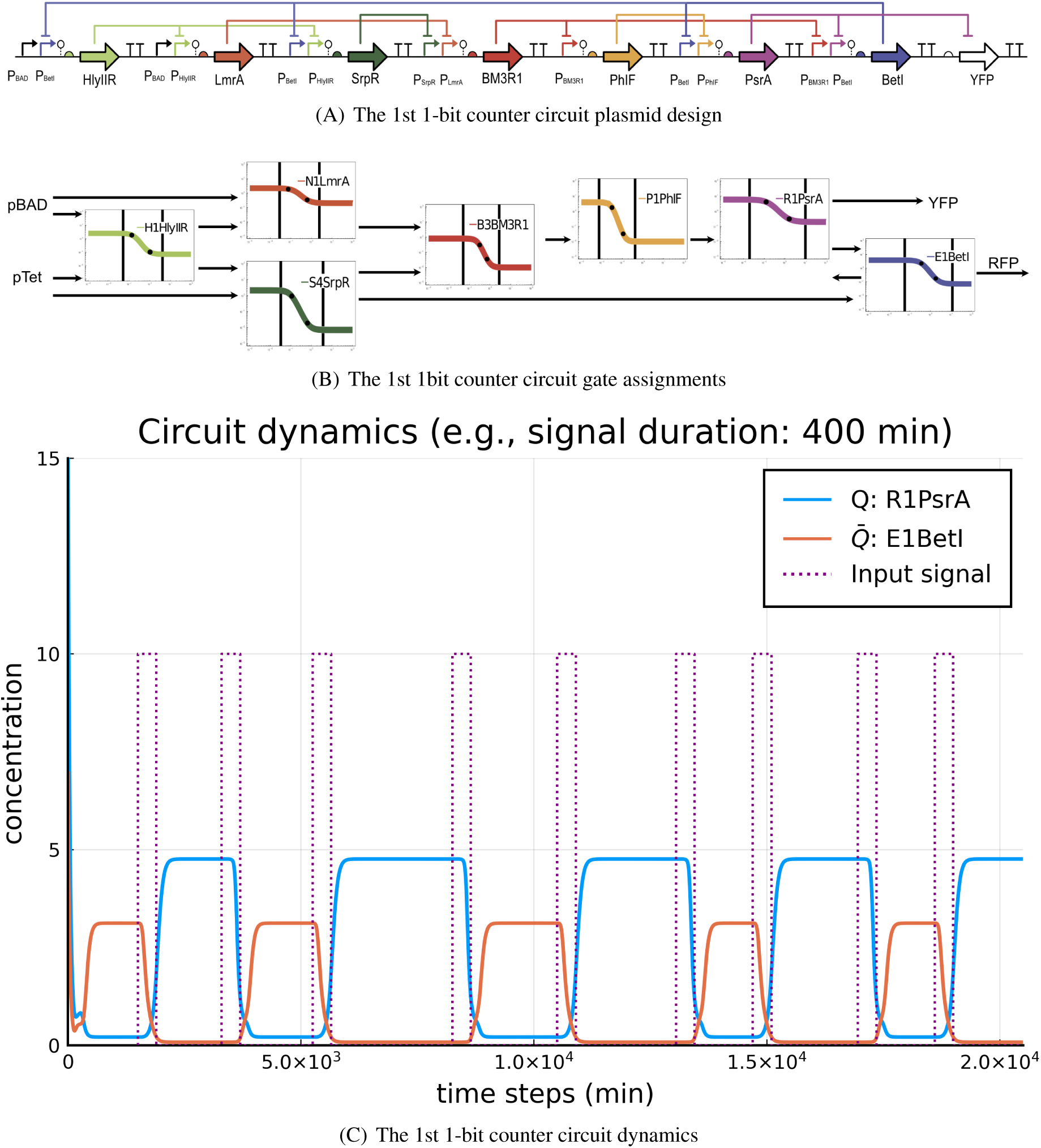
Related to Figure 2 & 15. The first 1-bit counter circuit with 7 unique experimental gates. This specific design is the output from the Cello 2.0 pipeline. (A) The circuit plasmid design. (B) The gate assignments for the associated plasmid in (A). (C) An example of the counter dynamics with *δ* = 400.

**Supplementary Figure 8:**
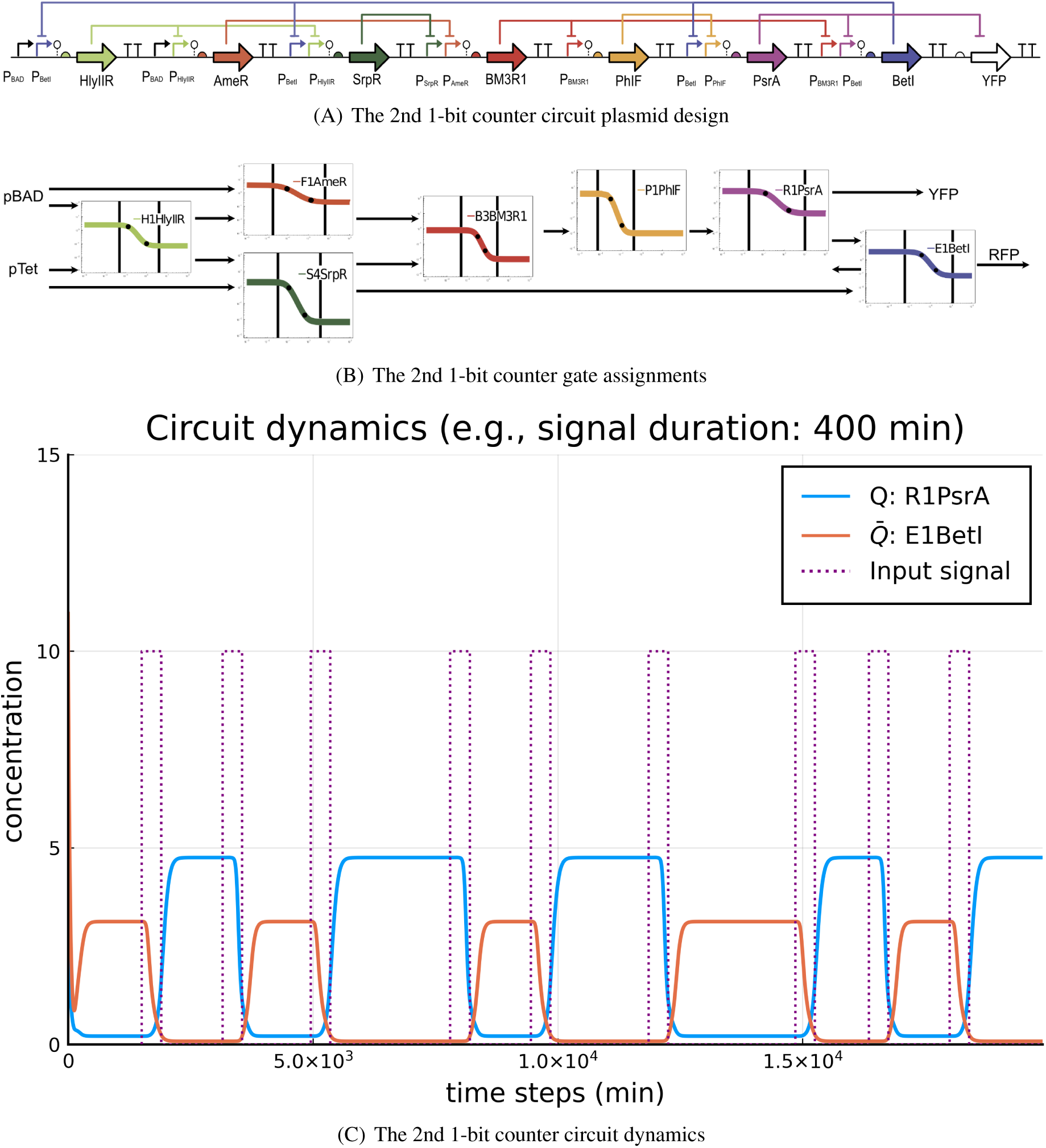
Related to Figure 2 & 15. The second 1-bit counter circuit with seven unique experimental gates. (A) The circuit plasmid design. (B) The gate assignments for the associated plasmid in (A). (C) An example of the counter dynamics with *δ* = 400.

**Supplementary Figure 9:**
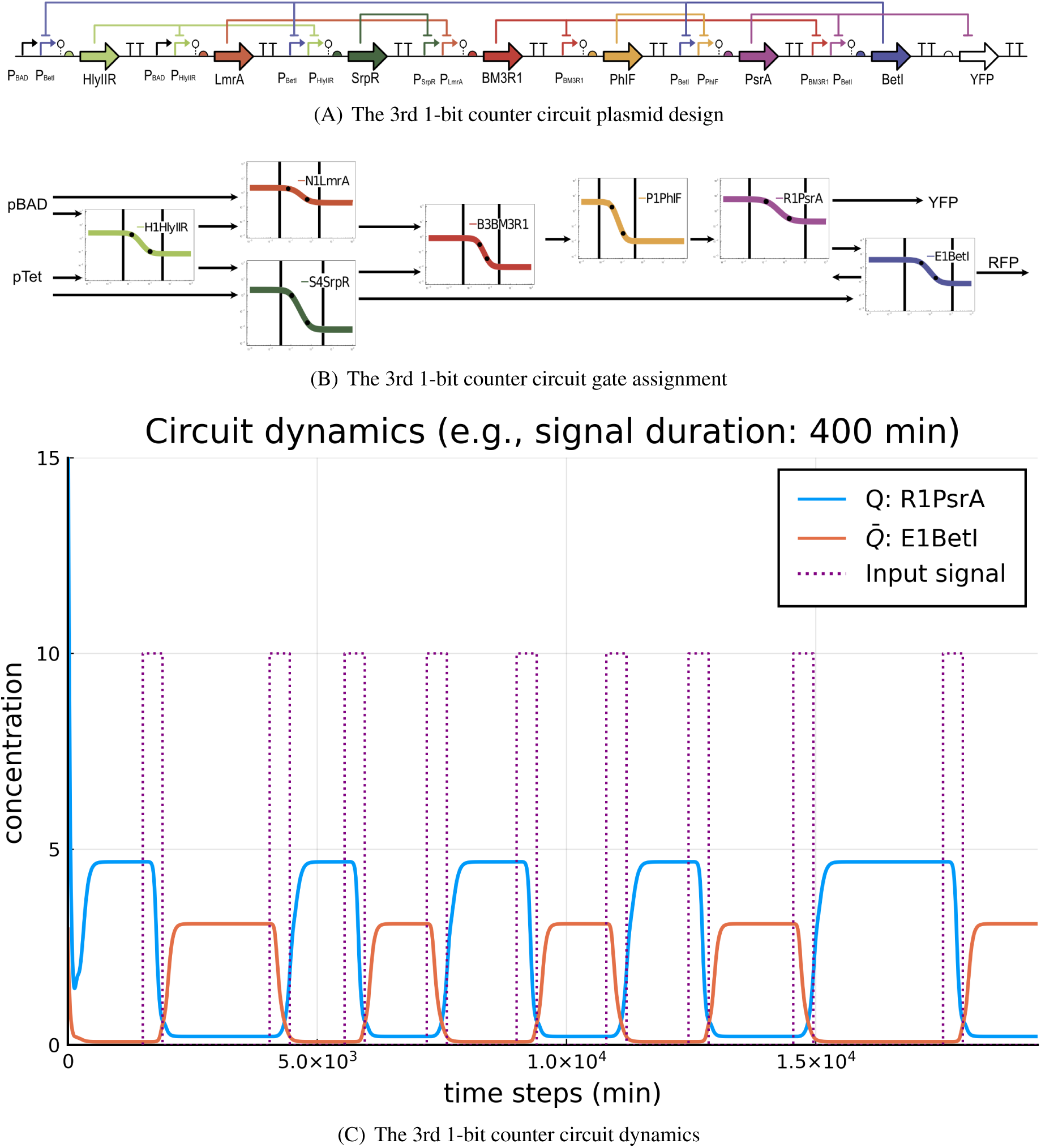
Related to Figure 2 & 15. The third 1-bit counter circuit with seven unique experimental gates. (A) The circuit plasmid design. (B) The gate assignments for the associated plasmid in (A). (C) An example of the counter dynamics with *δ* = 400.

**Supplementary Figure 10:**
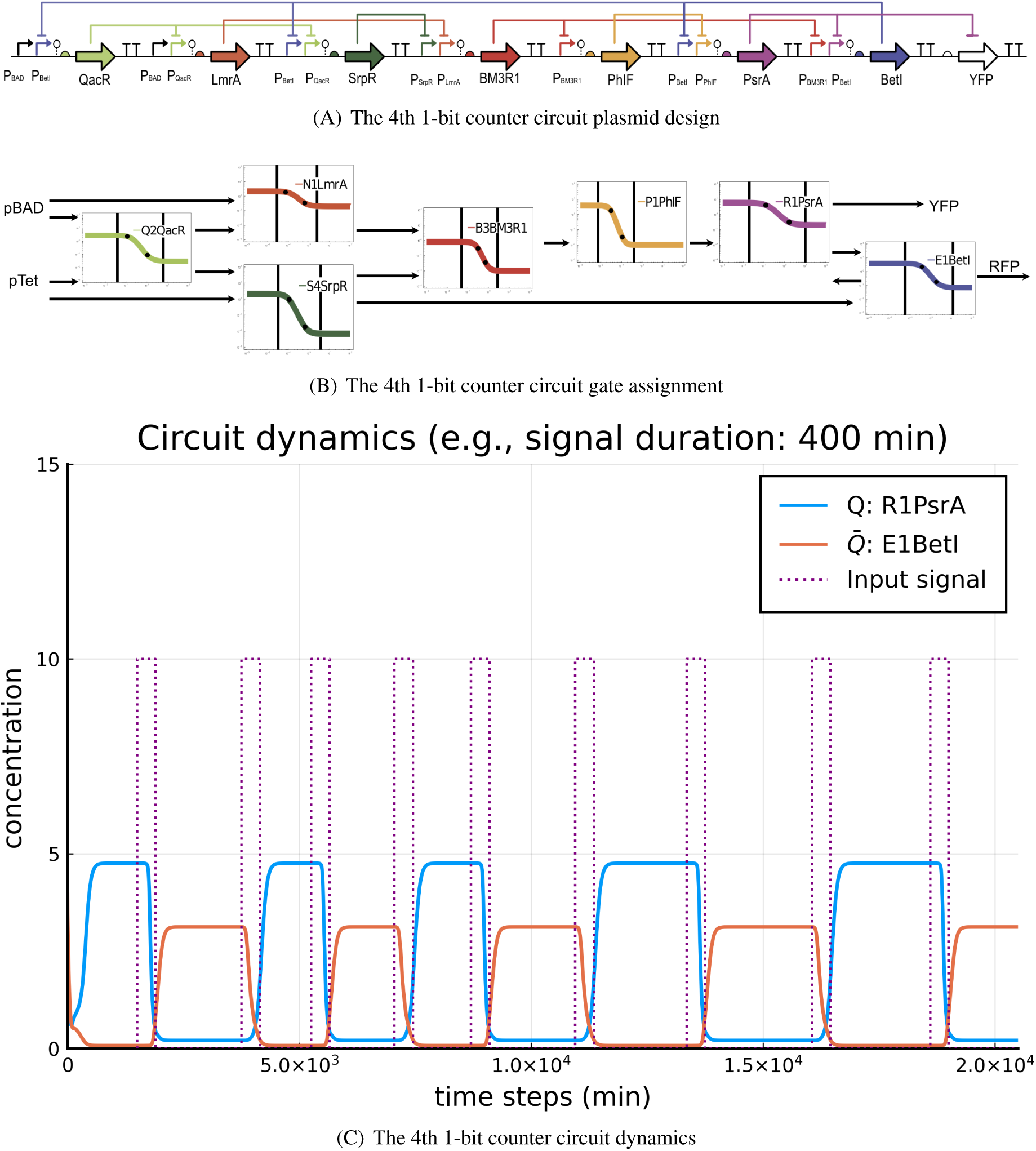
Related to Figure 2 & 15. The fourth 1-bit counter circuit with seven unique experimental gates. (A) The circuit plasmid design. (B) The gate assignments for the associated plasmid in (A). (C) An example of the counter dynamics with *δ* = 400.

**Supplementary Figure 11:**
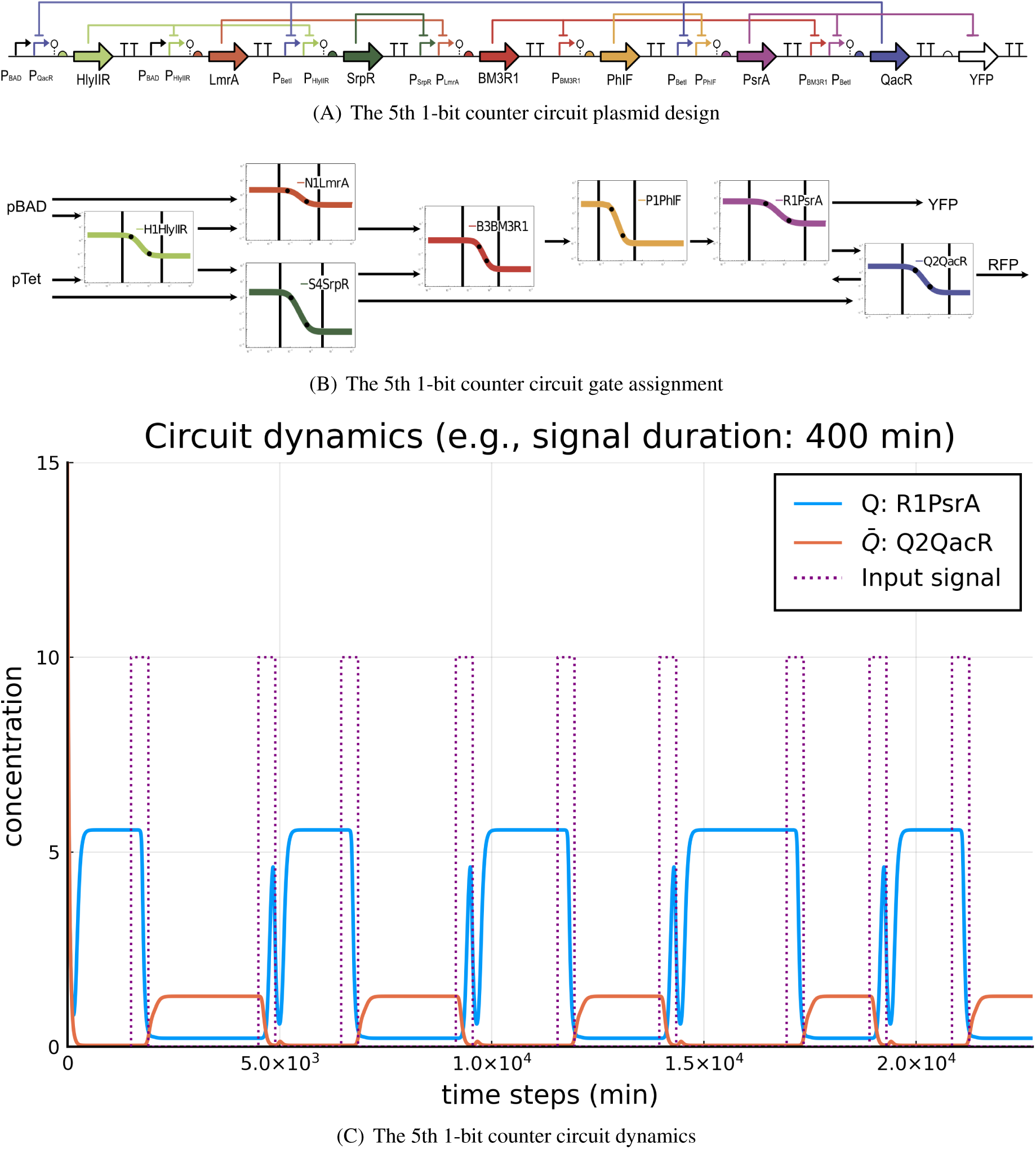
Related to Figure 2 & 15. The fifth 1-bit counter circuit with seven unique experimental gates. (A) The circuit plasmid design. (B) The gate assignments for the associated plasmid in (A). (C) An example of the counter dynamics with *δ* = 400.

**Supplementary Figure 12:**
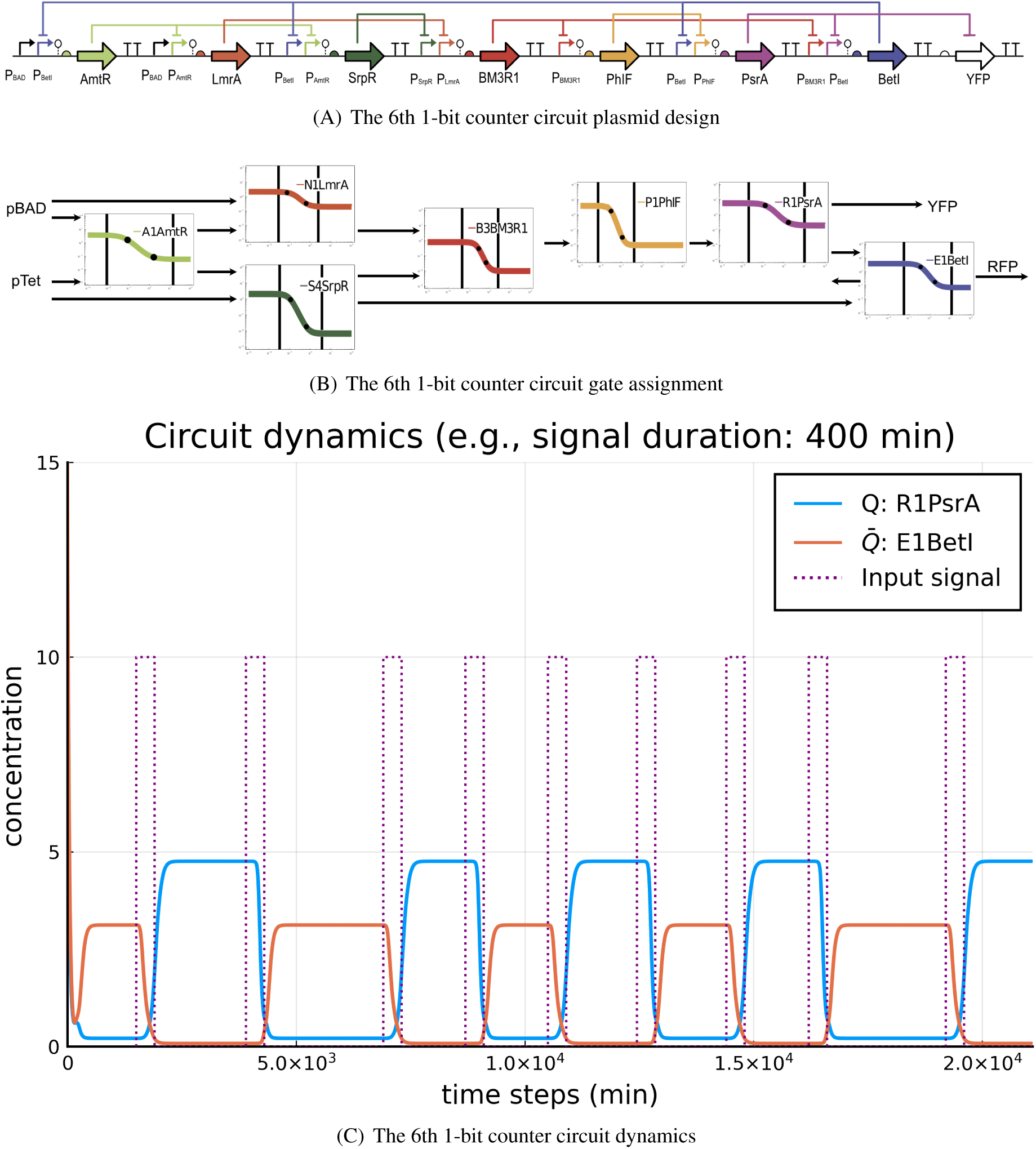
Related to Figure 2 & 15. The sixth 1-bit counter circuit with seven unique experimental gates. (A) The circuit plasmid design. (B) The gate assignments for the associated plasmid in (A). (C) An example of the counter dynamics with *δ* = 400.

**Supplementary Figure 13:**
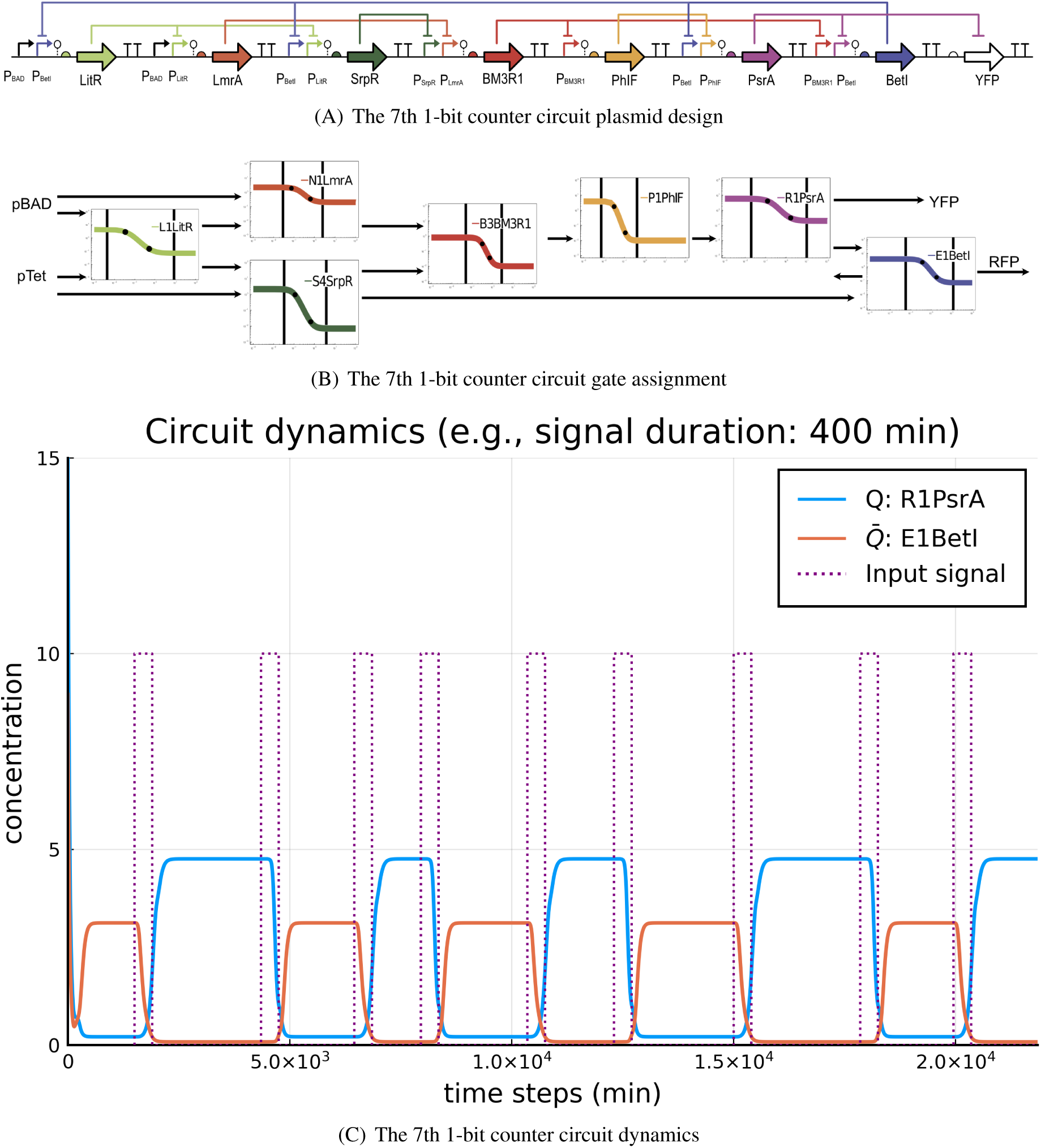
Related to Figure 2 & 15. The seventh 1-bit counter circuit with seven unique experimental gates. (A) The circuit plasmid design. (B) The gate assignments for the associated plasmid in (A). (C) An example of the counter dynamics with *δ* = 400.

**Supplementary Figure 14:**
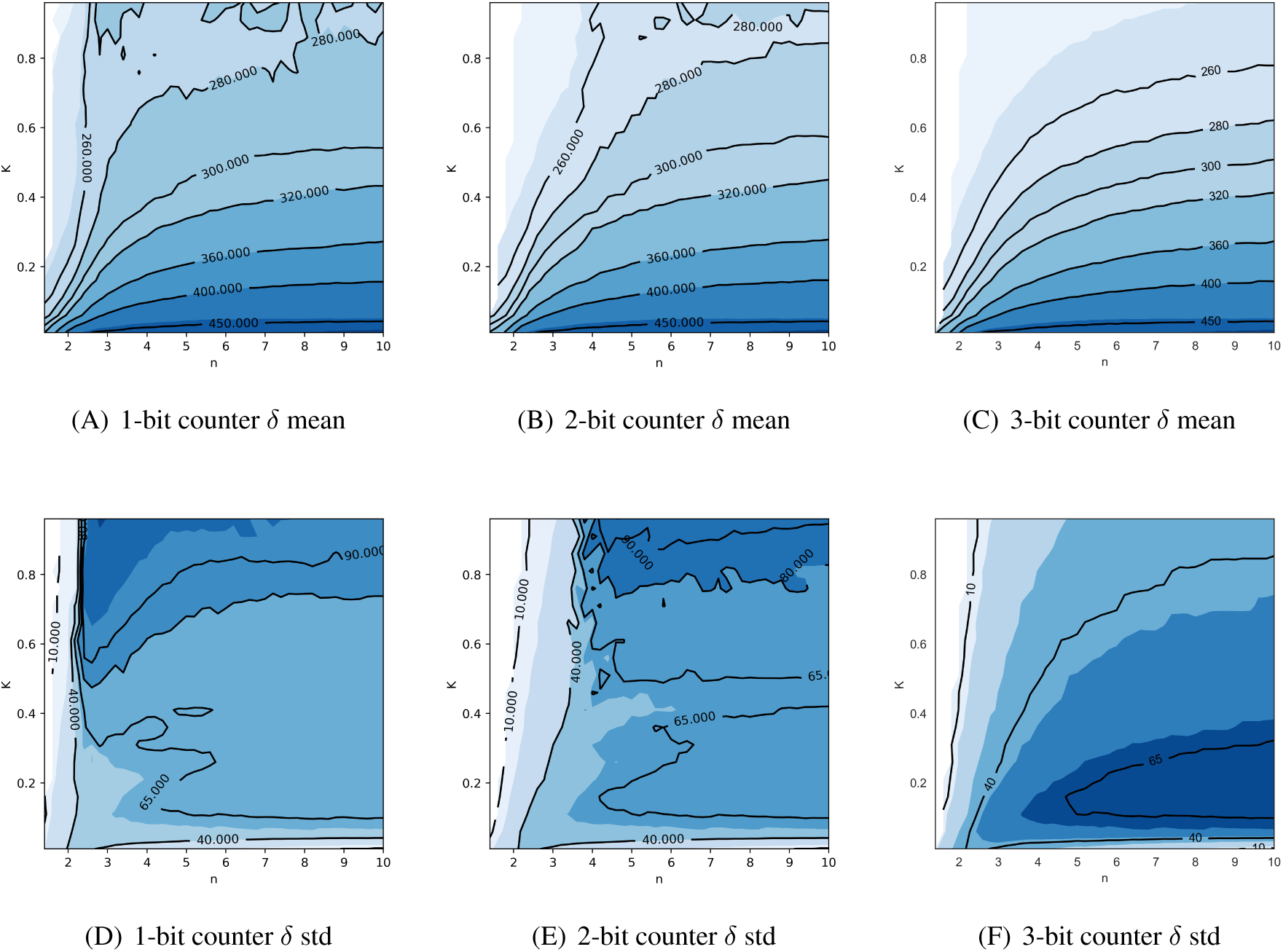
Contour plots of *δ*-mean and *δ*-std comparing between 1-bit, 2-bit, and 3-bit counters. Related to Figure 12 & 13. The 1st row shows the plots for *δ*-mean, and the 2nd row shows the plots for *δ*-std.

## Notes

### Competing Interest Statement

The authors have declared no competing interest.

